# Reward influences movement vigor through multiple motor cortical mechanisms

**DOI:** 10.1101/2025.04.09.648001

**Authors:** Adam L. Smoulder, Patrick J. Marino, Emily R. Oby, Sam E. Snyder, Aaron P. Batista, Steven M. Chase

## Abstract

The prospect of greater rewards often invigorates movements. What neural mechanisms support this increase of movement vigor for greater rewards? We had three rhesus monkeys perform reaching movements to targets worth different magnitudes of reward. We recorded neural population activity from primary motor and dorsal premotor cortex, brain areas at the output of cortical processing for voluntary movements, and asked how neural activity mediated the translation of reward into increased vigor. We identified features of neural activity during movement preparation, initiation, and execution that were both correlated with vigor and modulated by reward. We also found that the neural metrics that correlate with different aspects of movement vigor exhibit only limited correlation with one another, suggesting that there are multiple mechanisms through which reward modulates vigor. Finally, we note that the majority of reward’s modulation of motor cortical activity cannot be accounted for by reward-mediated vigor differences in behavior, indicating that reward modulations within motor cortex may serve roles in addition to affecting vigor. Overall, our results provide insight into the neural mechanisms that link reward-driven motivation to the modulation of the details of movement.

## Introduction

We only perform actions because we have sufficient motivation to do so. But motivation not only drives the binary decision to act or not, it also influences *how* we perform those actions. We try harder when more is on the line, whether it’s practice versus a playoff game, or traversing a wide walkway versus a narrow bridge. In an experimental setting, motivation is often manipulated by randomizing the reward to be received for successful performance on a particular trial and cueing each trial’s reward value in advance. An observation that has emerged from many of these experiments is that the prospect of greater reward often increases movement vigor^1^, commonly measured through decreases in reaction time^2–16^ or increases in movement speed^2,3,5–10,12,13,17–27^.

Here, we consider the possibility that the primary and dorsal premotor cortex (jointly referred to here as “motor cortex”) may play a critical role in the translation of reward into vigor. There are several reasons why we suspect this may be the case. First, motor cortex is the final cortical area that processes volitional movement commands before they are sent to the body^28^. Second, nonhuman primate studies of reaching have identified aspects of motor cortical activity that correlate with reaction time and peak speed during movement preparation^29–35^, initiation^36–40^, and execution^40–45^. Third, motor cortical activity is modulated by rewards at stake for a given task^14,35,46–55^. However, prior studies of reward-mediated vigor have focused primarily on subcortical brain regions^2,3,56–58^, and the role of motor cortex in translating increased rewards into invigorated movement is not well studied.

To study the relationship between reward, motor cortical activity, and movement vigor, we trained three monkeys on a delayed reaching task where the reward size for successful completion of a trial was pre-cued. We observed that vigor increased when greater rewards were made available. We then studied motor cortex activity during movement preparation, initiation, and execution, identifying correlates of movement vigor within each epoch that were also modulated by reward. We find that these motor cortical correlates of vigor only exhibit limited correlation with one another, supporting the argument that reward is affecting multiple distinct mechanisms that feed into vigor computations. Finally, we demonstrate that vigor does not account for the majority of reward’s effects on motor cortical activity. Overall, this work elucidates the nature of reward-driven motivational signals in motor cortex and their relationship with movement vigor.

## Results

We trained 3 rhesus monkeys to perform a challenging reaching task (**Fig. 1A**, see Methods for details)^32^. We tracked the hand’s position as the arm moved freely in space to control a cursor on a screen in front of the animal. After acquiring a central target to initiate a trial, a reach target appeared in one of multiple possible locations (Monkeys E and R: 8 locations, P: 4 locations), and either the color (Monkeys E and P) or image (Monkey R) inside the target indicated the reward to be received if the trial were performed correctly. We analyze trials across three reward sizes: Small, Medium, and Large (Methods). After a variable-length delay period, the center target disappeared, serving as a go cue. The animals then had to quickly move the cursor into the target with an arm reach and hold the cursor in the target for 400 ms to receive the cued reward. The task was challenging in terms of both its speed and accuracy requirements, requiring brisk reaches to small targets. As a result, success rates were approximately 80% (see **Fig. S1** for failure mode breakdown).

**Figure 1.**
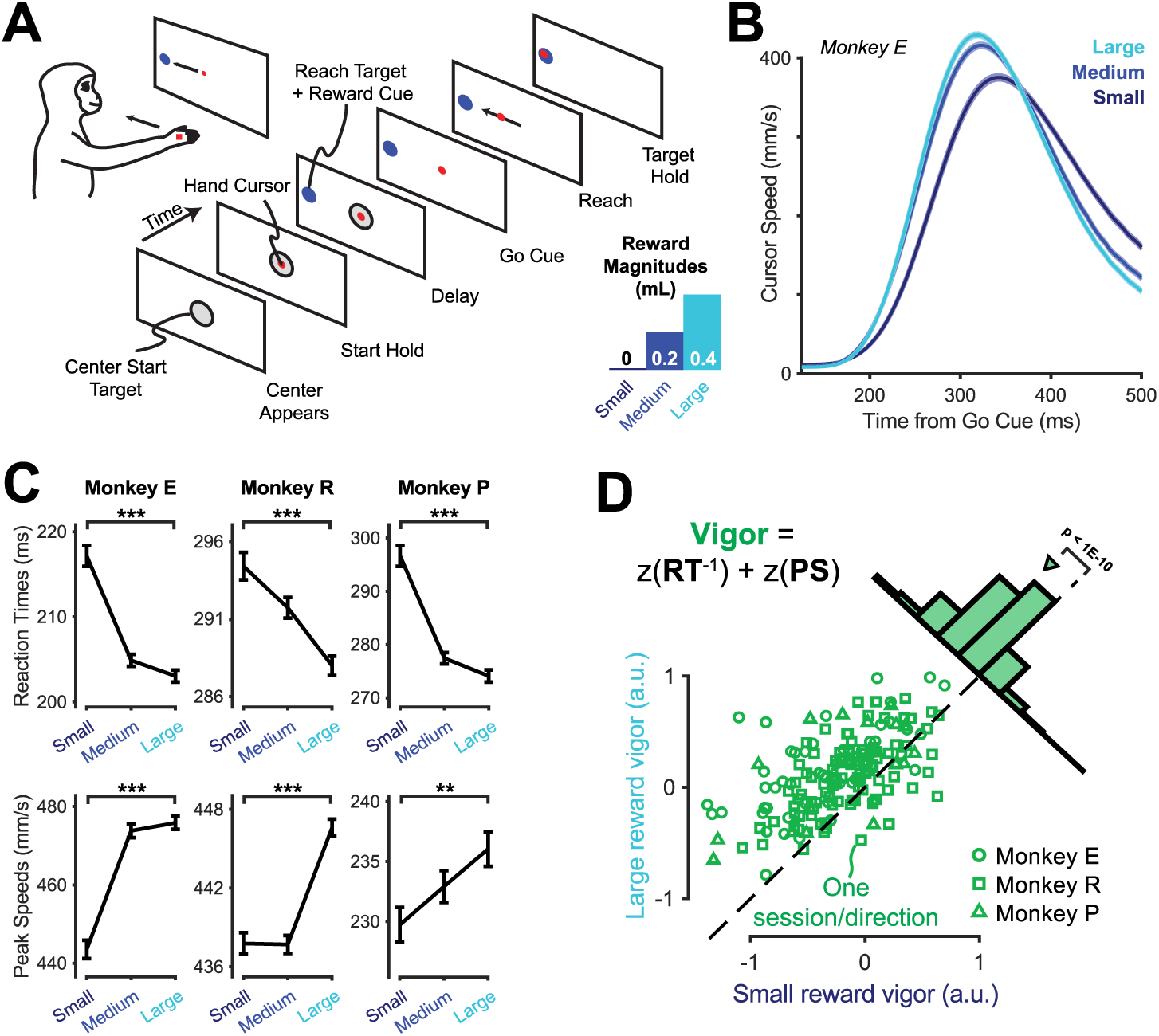
Reward magnitude affects movement vigor in a reaching task. (A) We trained monkeys to perform a delayed reaching task where the image within the reach target cued the amount of liquid reward at stake for the trial (see Methods for task details; reward sizes for Monkey E shown). Colors here (Small = dark blue, Medium = blue, Large = cyan) are for visualization and are not the actual cues used. Reward sizes for the other animals were: Monkey R - 0, 0.22, 0.44 mL, Monkey P - 0.15, 0.3, 0.6 mL. (B) Higher reward trials show shorter latency to movement and greater movement speed. We show the average cursor speed profile from Monkey E over time split by reward condition (mean ± S.E. shading, though S.E. is small and hard to see). (C) Greater reward speeds leads to both faster reaction time (top; median ± S.E., ***p < 0.001, Wilcoxson rank-sum test) and peak movement speed (bottom; mean ± S.E., **p < 0.01, ***p < 0.001, Welch’s t-test). See **Table S1** for all main figure statistics. (D) Reward improves vigor. We show mean vigor for Large versus Small reward trials for each animal-direction-session. We define vigor in the equation shown at the top-left, where RT is reaction time, PS is peak speed, and z(x) is the z-score operation (Methods). The inset shows a histogram of differences in vigor: 134 out of the 162 direction x day conditions (83%) showed greater vigor for Large reward trials compared to Small trials (median = 0.31, sign-rank test p < 10^-10^).

### Reward improves movement vigor

We first wanted to determine if reward modulates reaching vigor in this task. We capture the vigor of a movement in terms of its reaction time and peak speed (Methods). The two metrics were significantly correlated, albeit weakly (median Spearman rank correlation r = -0.21, p < 0.001, sign-rank test, **Fig. S1**). For each animal, we found that reward modulated vigor: comparing Large to Small reward, all three animals exhibited reductions in reaction time and increases in peak speed (**Fig. 1B-C**). Previous studies of movement vigor have observed effects on either or both of these metrics. For this work, we define “vigor” by combining reaction time and peak speed, giving equal weight to each:

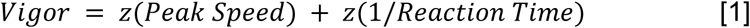

Where *z(x)* is the z-score operation across all trials of all conditions within each animal (Methods). We invert reaction time both so that a higher value means greater vigor and also because it results in data that align better with a normal distribution^59,60^. We found that reward significantly increased movement vigor (**Fig. 1D**).

While the animals performed the task, we used implanted multielectrode Utah arrays to record neural population spiking activity from primary motor cortex and/or dorsal premotor cortex (Methods), which we collectively refer to here as simply “motor cortex”. Given that we observe invigoration of movements with reward, we can now step through the epochs of the task and examine motor cortex activity, comparing the effects of cued reward with the animals’ movement vigor on each trial.

### Reward modulates preparatory neural activity correlated with vigor

We begin by investigating movement planning. Each trial began when the animal held the cursor in the center start location. This triggered target onset, where a target appeared that specified both the reach goal and the volume of reward to be received for a successful reach. Previous work in macaque reaching tasks has demonstrated robust evidence of movement preparation in motor and premotor cortex spiking activity during the delay period that follows target onset, as the animal waits for the go cue^30,34,61,62^. Here we investigate links between vigor and reward modulation in this preparatory activity.

To examine neural activity during movement preparation, we first made peri-stimulus time histograms (PSTHs) of motor cortical single unit activity as a function of reward. We observed clear differences between Small and Large reward trials (**Fig. 2A**). We next made PSTHs as a function of the vigor in the upcoming reach. To construct these, we used only Medium reward trials and split them up into Low vs. High vigor based on the vigor of the trial’s movement (Methods). Similar to reward, we saw vigor-related differences in neural activity (**Fig. 2B**). We found that the tuning of individual neurons for reward and vigor was correlated (see **Fig. S2A** for quantification). For example, units with higher firing rates for a Large reward trial compared to a Small reward trial also tended to have higher firing rates for an upcoming High vigor trial compared to a Low vigor trial (e.g., units 299 and 66). We also saw that the vigor tuning was consistent across reward sizes: the tuning to vigor within Small and Large trials was highly similar to that of Medium rewards (**Fig. S2B-C**).

**Figure 2.**
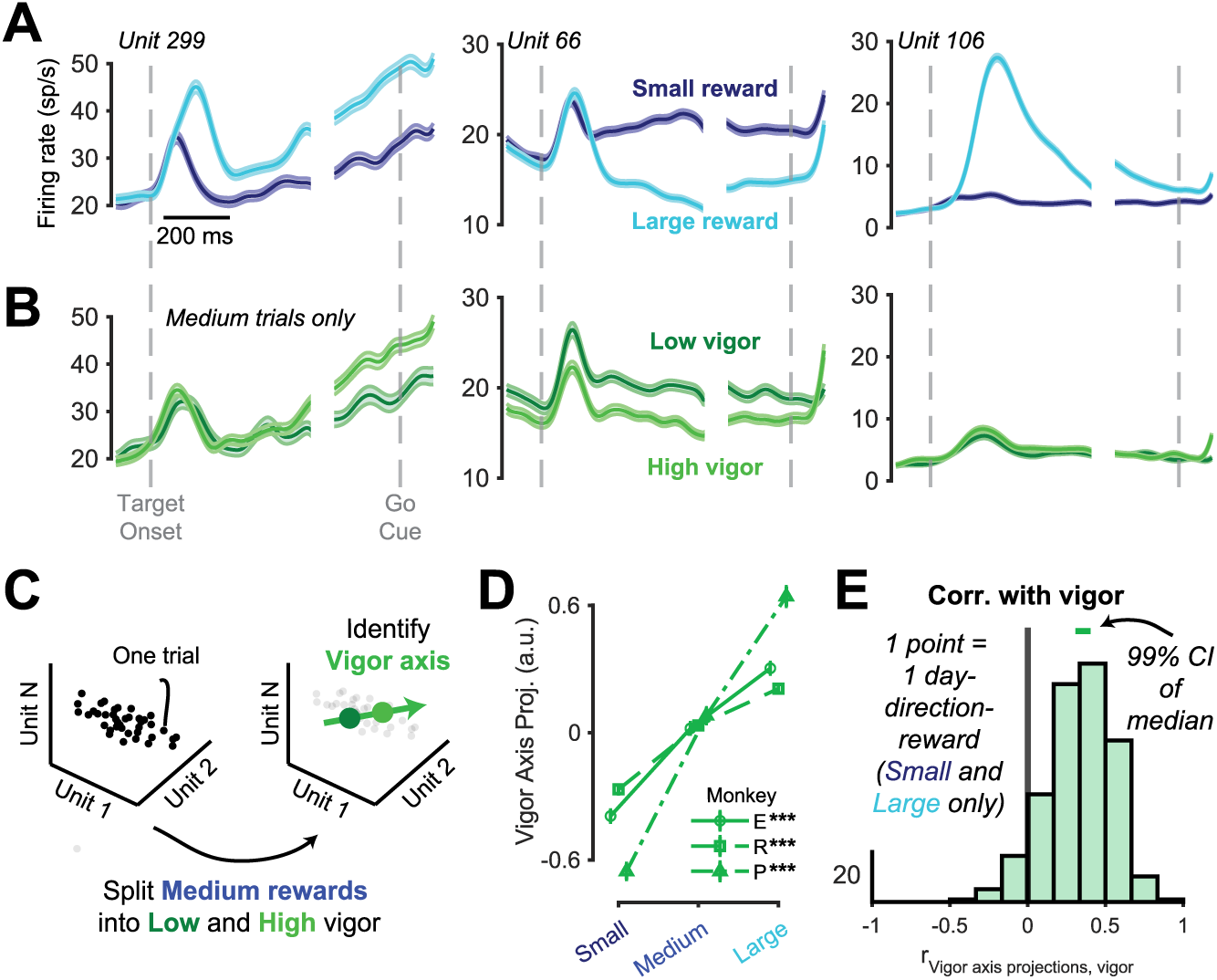
Reward modulates vigor-related patterns of motor cortical preparatory activity. (A) Motor cortex exhibits firing rate changes in response to different cued rewards. We show PSTHs (mean ± S.E.) from 3 example units from Monkey E as a function of reward cue. For visual simplicity, we only show Small (dark blue) and Large (light blue) reward trials. (B) Motor cortex shows firing rate differences encoding the vigor of the upcoming movement. We took Medium reward trials and split the data in half (based on the median vigor) into Low vigor (dark green) and High vigor (light green) trials, then averaged across these to make PSTHs. (C) We identified a “vigor axis” in the neural population state space as a dimension that separated High from Low vigor trial averages (large points) within Medium reward trials (small points) at the end of movement preparation, (-150 ms before go cue through 50 ms after). We note that we considered alternative ways of calculating the vigor axis and saw few differences in the ability of the neural activity to predict vigor (**Fig. S3**). (D) Vigor-axis projections are modulated by reward. We calculated the mean (± S.E. error bars, which are present but small) as a function of reward after z-scoring values within each day-direction condition. Stars indicate statistically significant difference in means between Small and Large reward conditions for each animal (***p < 0.001, Welch’s t-test). (E) Vigor-axis projections are correlated with vigor in held-out conditions. We calculated the Spearman rank correlation within each day-direction-reward condition that had at least 10 trials (n = 318) and made a histogram of the correlation values (median = 0.35). Because the vigor axis was found using Medium reward trials, we only use Small and Large trials for this analysis. We show a 99% confidence interval of the median value in a bar at the top.

We can more holistically assess vigor and reward effects by studying them at the neural population level. To do this we treat the firing rate of each neuron as a dimension in a high-D space^63^. In this space we can correlate modulations of neural activity that are informative about metrics or task parameters of interest, such as vigor and reward, to see how they relate to one another. An advantage of this approach is that it allows us to “stitch” neural recordings across multiple recording sessions using factor analysis^64–66^, improving statistical power in assessing potentially weak effects of reward or vigor (see Methods). For the remainder of this work, we perform analyses on the factors scores found from this neural stitching procedure (Monkey E: 24 factors, R: 18 factors, P: 16 factors).

We identified a “vigor axis” in the neural population activity across all reach direction conditions to capture the change in the activity of the neural population that correlated with vigor (**Fig. 2C**, Methods). To do this, we took a bin of neural activity at the end of the delay epoch (from 150 ms before the go cue to 50 ms after) to capture the end of movement preparation. We then split Medium reward trials into Low and High vigor trials based on the median vigor, and we identified the vector in the neural state space that connected the average neural activity for the Low vigor trials with that of the High vigor trials. This gave us the “vigor axis”.

To assess how reward might drive vigor changes, we then projected neural activity from trials from all *reward* conditions along this vigor axis. If the reward signals in motor cortex serve to invigorate actions, then we would expect that the neural metrics correlated with vigor are similarly increased by reward. To test this for the case of the vigor axis that we identified, we calculated the average vigor-axis projection within each reward condition. We found greater rewards indeed yielded greater vigor-axis projections (**Fig. 2D**).

We wanted to know how well vigor axis projections correlated with vigor within different task conditions (e.g., different rewards, different reach directions). We calculated the correlation between vigor-axis projections and (behavioral) vigor values within each day-direction-reward condition (Methods). We found that the two were substantially correlated even within Small and Large reward trials, which were not used in the vigor axis fitting process (**Fig. 2E**). The intersection of these two analyses demonstrates that vigor-axis projections are both (1) modulated by reward, and (2) correlated with movement vigor even when reward and other task conditions are held constant. This supports the hypothesis that reward is modulating patterns of motor cortex preparatory activity that are conducive to generating greater movement vigor.

As an additional analysis, we used reaction time or peak speed separately to identify axes that predicted these parameters alone. We found that the reaction time axis and peak speed axis were effectively the same as the vigor axis (**Fig. S3**). This implies that across-direction reaction time and peak speed encoding are effectively the same in motor cortex’s preparatory activity, lending credence to using a combined vigor metric as we have done.

Signals in motor cortex related to directional tuning have also been documented to correlate with upcoming movement vigor^34^ and to be impacted by reward^35^. To dissect this relationship we first estimated the directional tuning by identifying a 2D “target plane” in neural population activity that optimally separated the different reach directions (**Fig. 3A-B**, Methods). We then projected all trials into this target plane. For visualization, we calculated the average neural activity in the target plane for Small and Large reward sizes (**Fig. 3C**). We can see that increasing reward magnitude pushes neural activity further apart in the target plane, which may support better task performance^35^. Then, to look at vigor, we took the Medium reward trials and split them into Low and High vigor (**Fig. 3D**). Like reward magnitude, increasing vigor also improves target discriminability in this target plane.

**Figure 3.**
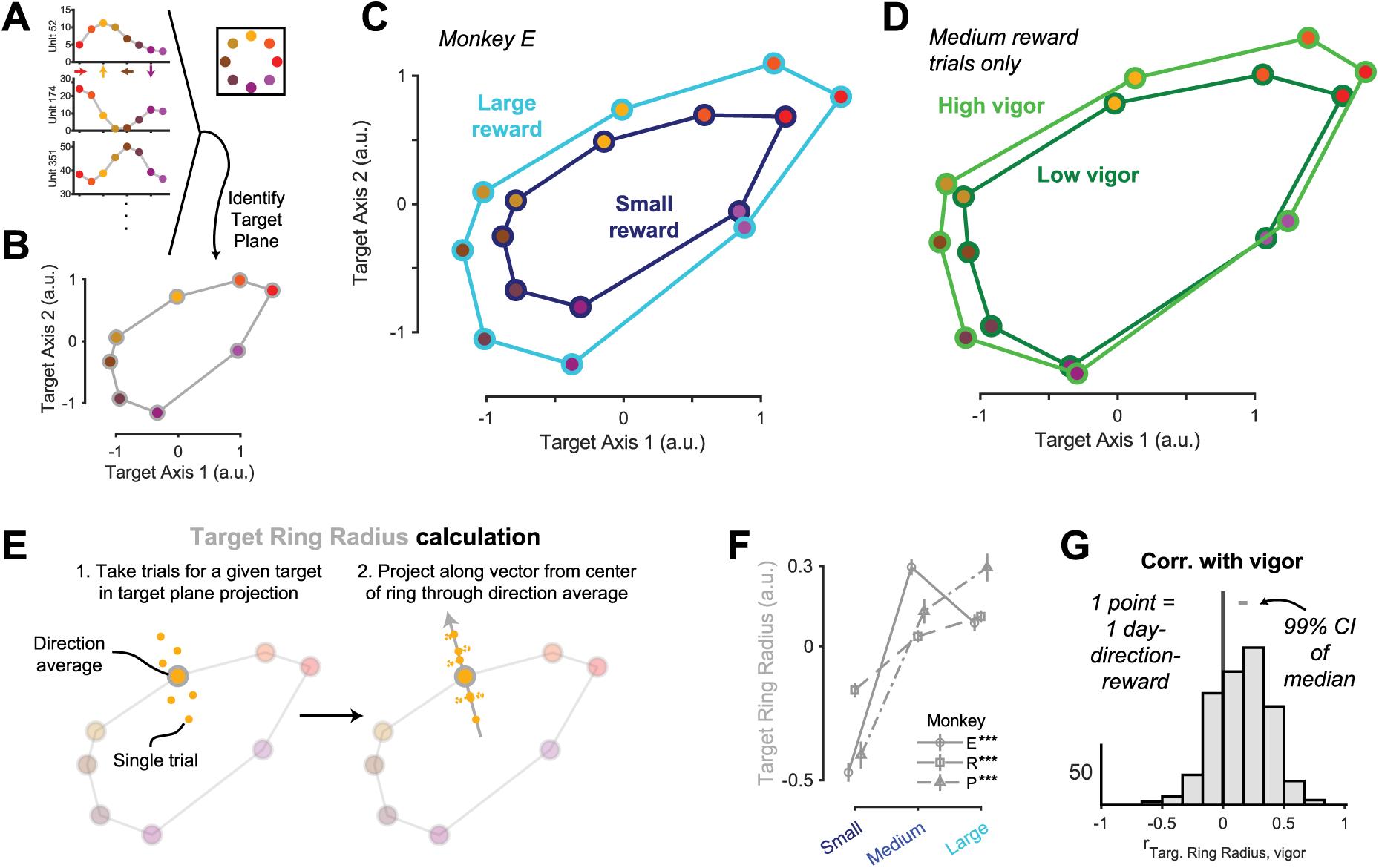
Greater reward and greater vigor both improve encoding of upcoming reach direction. (A) Neurons in motor cortex showed directional tuning during movement preparation, evidenced by these example tuning curves from Monkey E. A color legend for different reach directions is shown in the rosette. (B) We identified a “target plane” using linear discriminant analysis (LDA) to maximize separability between the direction conditions (Methods). Shown here is the average activity for each reach direction (dots, color indicates direction) and lines connect adjacent reach targets in a ring (gray) for visualization. (C) Neural preparatory states for different reach directions are spread farther apart for Large reward trials compared to Small. We show the average activity for each reach direction in Monkey E’s target plane in the same format as panel B for Small (dark blue ring) and Large (light blue ring) trials. (D) Neural preparatory states for different reach directions are spread farther apart for High vigor trials compared to Low. Same format as panel C, but using the Medium reward data split by Low (dark green) and High (light green) vigor. (E) We quantified the target ring radius on a single trial basis (see Methods for details). We took all trials’ projections into the target plane within a given reach direction (upward target trials shown as small dots in this example). We then drew a vector from the center of the target ring through the average for that reach direction and projected all trials along it. Dots schematize single-trial responses. (F) Reward modulates target ring radius in motor cortical preparatory activity. Same format as Fig. 2D (***p < 0.001 Small to Large reward difference, Welch’s t-test). (G) Target ring radius correlates with upcoming movement vigor within day-direction-reward conditions (n = 476). Same format as Fig. 2E, though all reward conditions are included (median correlation = 0.16).

To quantify this effect, we calculated a “target ring radius”^35^ value for each trial (**Fig. 3E**, Methods).We found that the target ring radius increases from Small to Large rewards for all animals (**Fig. 3F**) and that it weakly-but-significantly correlates with vigor on a trial-by-trial basis (**Fig. 3G**). Thus, in both target-agnostic (Fig. 2) and target-specific (Fig. 3) signals, we find that reward affects patterns of neural preparatory activity in a way that aligns with greater upcoming movement vigor.

### Reward accelerates both the latency and dynamics of movement initiation

We now progress from considering the movement preparation epoch to the movement initiation epoch. One of the strongest known neural correlates of movement vigor is the relationship between reaction time and the timing of neural “trigger” signals in the motor cortex. Trigger signals are changes in neural activity that occur irrespective of task condition before the start of an arm movement and are strongly predictive of reaction time^36,38,40,67^. We wondered whether aspects of this trigger signal might be affected by reward in ways that could modulate movement vigor.

To identify a trigger signal, we examined neural activity between the time of the go cue and movement onset. Single-unit activity reveals robust changes during this transition (**Fig. 4A**). To find an across-condition dimension that best captured these transition dynamics, we used linear discriminant analysis (LDA) to separate pre-movement activity from peri-movement activity (**Fig. 4B**; Methods). We define this dimension as the trigger axis.

**Figure 4.**
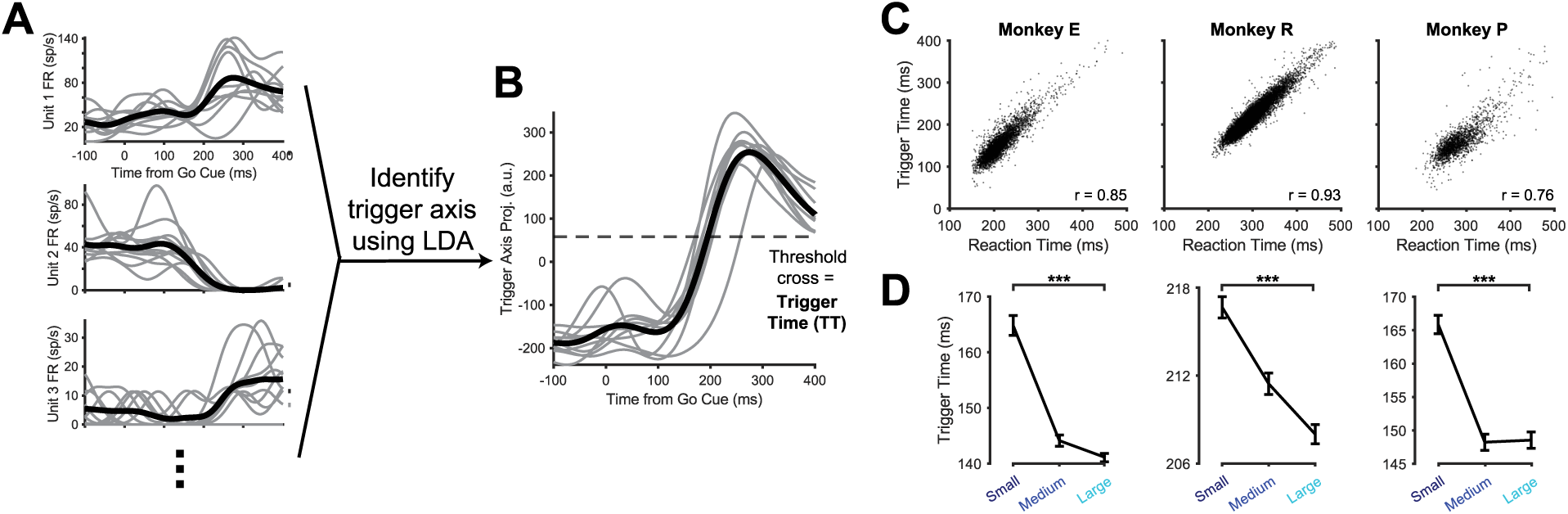
Reward modulates the timing of trigger signals in the motor cortex. (A) We examined firing rates at the transition period between movement preparation and execution, occurring between the time of the go cue and movement onset. Example units from Monkey E are shown, with single trials in light gray and the average across them in black. (B) We use LDA to identify an across-condition “trigger axis” that captures the motor cortical activity transition occurring preceding movement onset. We then identified a threshold value where the timing of single trial threshold crossing (trigger time) maximally correlated with reaction time. The same trials are shown here as in panel A. (C) Trigger time and reaction time are highly correlated. Spearman rank correlations are given in the bottom-right corner of each plot. (D) Trigger time decreases as a function of increased cued reward. The median (± S.E.) of the trigger time is shown for each reward condition, and statistics compare Small and Large reward trials (***p < 0.001, Wilcoxson rank-sum test).

After establishing a trigger axis, we used it to calculate the time of a neural threshold crossing, or “trigger time” (Methods). To do this, we projected neural activity during single trials onto the trigger axis, producing a trial-by-trial trigger signal. We then computed the time at which the trigger signal crossed a threshold, which we called the “trigger time”. We selected the threshold value that maximized the correlation between the trigger time and the reaction time. The trigger time was highly predictive of reaction time (**Fig. 4C**), a reassuring finding that confirms other reports of a close correlation between neural activity in motor cortex and movement reaction time^36,68^. We then examined how reward affected the trigger time. We observe that, much like reaction time, greater reward corresponded to a faster trigger time (**Fig. 4D**).

How might reward modulate the neural trigger process to achieve a faster trigger time? If we fix a static threshold across conditions, we can identify 3 factors that could affect the trigger time (**Fig. 5A**, see Methods for calculation details). First, the time at which the signal starts to change could occur earlier. We call the time at which the signal starts changing the changepoint time. Second, the speed of the neural dynamics (i.e., the slope of the transition from the changepoint to the trigger time) could increase. We call this the trigger speed. Finally, the initial neural distance from threshold could decrease, allowing the trigger time to be hit earlier. We call this the initial distance. Each of these metrics reflects different aspects of neural activity: the initial distance reflects the neural state when the animal begins initiating the movement, the trigger speed reflects the gain on neural dynamics, and the changepoint time relates to how quickly a trigger command arrives to motor cortex. We evaluated each in the same manner as the neural metrics during movement preparation, seeking to find not only which aspects of the transition were affected by reward, but which related to vigor on a trial-by-trial basis.

**Figure 5.**
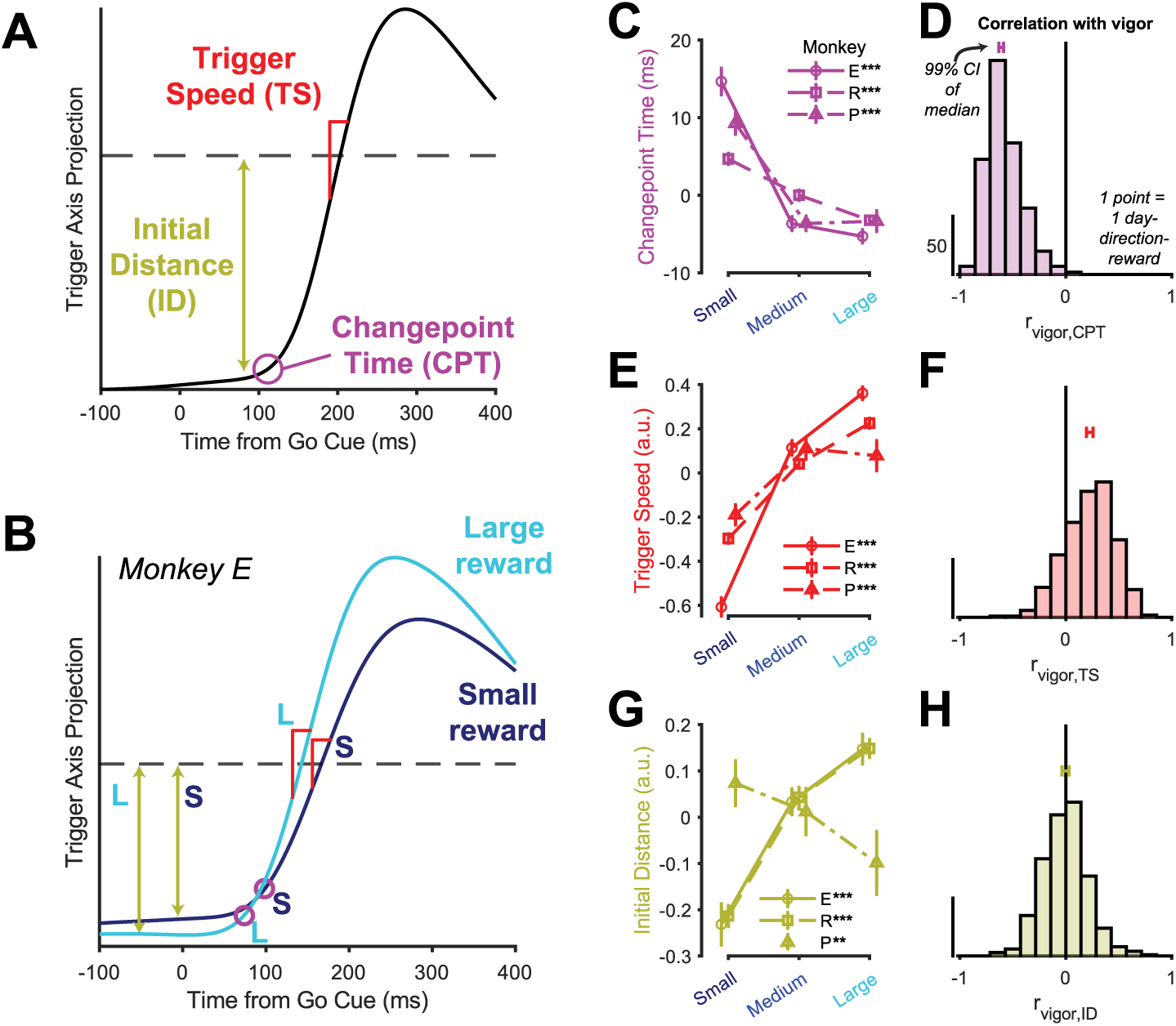
Reward modulates multiple aspects of trigger signals related to vigor. (A) Given a trigger threshold and a single trial trigger axis trajectory, we identify three aspects of movement initiation that have the potential to determine trigger time: the changepoint time (CPT, purple), the trigger speed (TS, red), and the initial distance (ID, gold). Any one or combination of these may be affected by reward to yield the earlier trigger time observed with greater reward seen in Figure 4. (B) Average Small and Large reward trigger axis projections for Monkey E. We illustrate the quantities identified in panel A for each reward size. (C) Greater reward yields faster changepoint times. We show changepoint time for each reward size (median ± S.E. error bars). For visualizing results across animals, we centered values on their overall median changepoint times (Monkey E = 87.5 ms, R = 151.5 ms, P = 99.8 ms). Statistical comparison between Small and Large rewards is shown for each animal in the legend (***p < 0.001, Wilcoxson rank-sum test). (D) Vigor is strongly correlated with changepoint time. Spearman rank correlations were calculated within each day-direction-reward condition that had at least 10 trials available (n = 489). We show a 99% confidence interval of the distribution’s median in a bar above the histogram (median r = 0.81). All individual animals’ statistics are listed in **Table S1**. (E) Reward modulates trigger speed. Same format as panel B, but using mean instead of median (**p < 0.01, ***p < 0.001, Welch’s t-test). (F) Trigger speed correlates with vigor (median r = 0.23). Same format as panel D. (G) Reward affects initial neural distance from threshold. Same format as panel E. (H) Initial distance shows little correlation with vigor (r = -0.00). Same format as panel D.

We found that two of the three trigger factors were influenced by reward in a manner conducive to improving vigor (**Fig. 5B**). First, we observed that changepoint time robustly decreased with reward (**Fig. 5C**) and was strongly correlated with vigor (**Fig. 5D**). Second, we saw that trigger speed strongly increased as a function of greater reward (**Fig. 5E**) and demonstrated a middling-but-robust correlation with vigor (**Fig. 5F**). Finally and most curiously, initial distance showed a strong *increasing* trend with reward for two of the three animals, which would predict a longer duration to passing the trigger threshold (**Fig. 5G**). However, we saw minimal correlation of initial distance with vigor (**Fig. 5H**), suggesting it plays a minor role in determining vigor. We note that the results shown here were broadly consistent across a variety of methods used to calculate the trigger axes (**Fig. S4A-C**), were significant using either reaction time or peak speed instead of the combined vigor metric for correlations (**Fig. S4D-E**), and that all highlighted trends were statistically significant within each animal (**Table S1**). These results indicate that reward accelerates both movement initiation time and neural trigger dynamics in a manner that corresponds to greater movement vigor.

### Like vigor, reward amplifies peri-movement neural dynamics

We next consider the movement epoch. Because peri-movement dynamics occur after the reaction time has taken place, here we focus on peak speed as our metric for vigor. Prior work has demonstrated that motor cortical activity during movements has two key parameters that have correlated with hand movement speed: (1) the magnitude of the neural trajectory^43,44^ which corresponds to the strength of motor cortical activation during movement, and (2) the speed at which the neural activity traverses this trajectory^40,45^, which reflects how quickly motor cortex is sending the sequence of movement signals down to the spinal cord. Reward could potentially be manipulating either (or both) of these to impact movement vigor.

To examine these effects, we performed principal components analysis (PCA) to identify dimensions that capture temporal dynamics irrespective of reach direction or reward conditions (Methods). We then visualized the average activity in the top two PCs as a function of reward and (for Medium reward trials) Low versus High peak speed. We qualitatively found that both greater reward and greater movement speed resulted in “stretched” neural trajectories (**Fig. 6A-B**).

**Figure 6.**
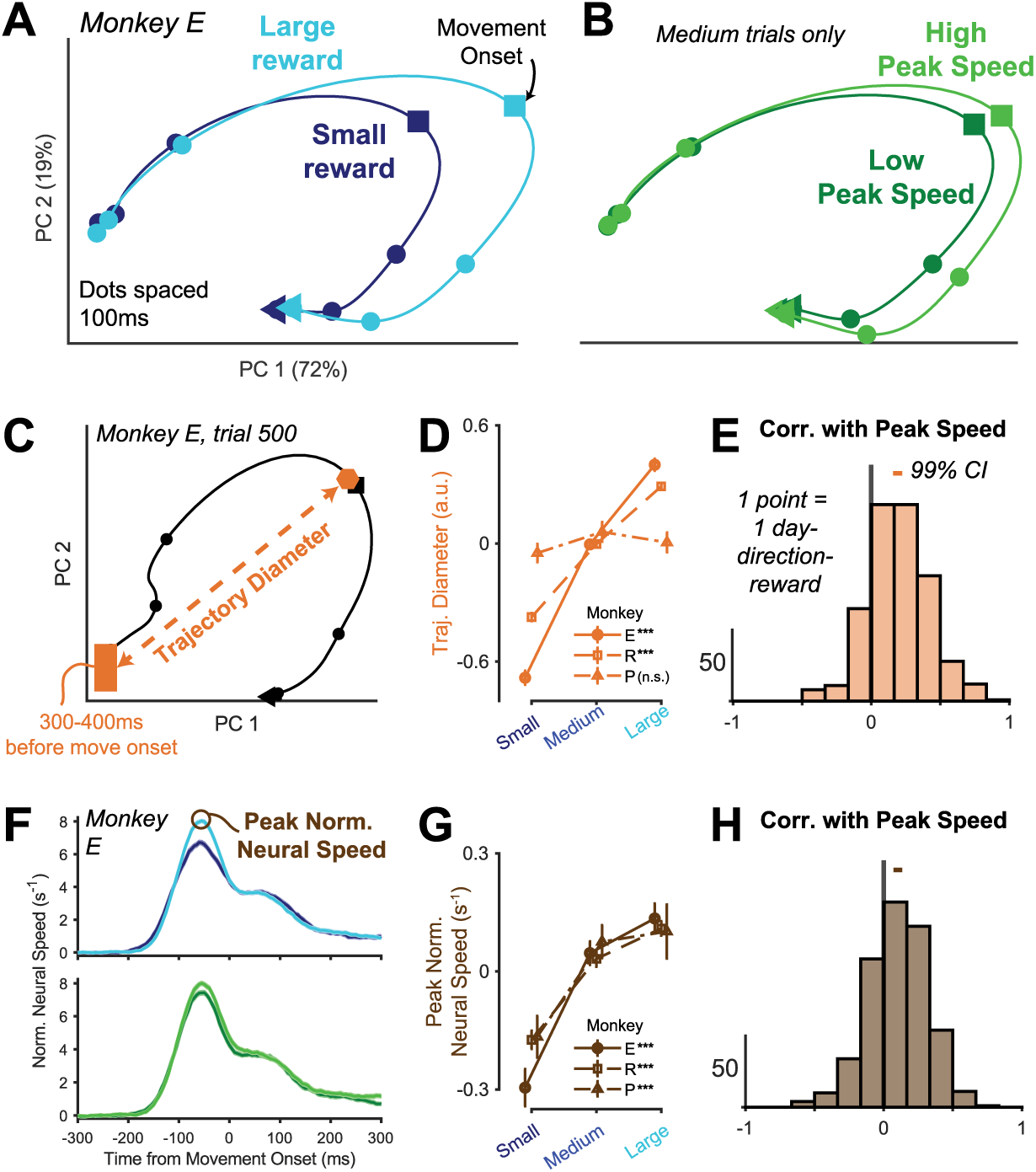
Reward both amplifies and quicken neural trajectories, corresponding to more vigorous movements. (A) Neural trajectories plotted in a 2D space that maximizes temporal variance (Methods). Trial-average trajectories are split by reward condition (Small = dark blue, Large = light blue). Percentages in the axis labels show the amount of variance in the average neural trajectory explained by each dimension (total of >90% for all animals). (B) Same as panel A, but for Low and High peak speed trials of the Medium reward condition only. (C) We quantified trajectory diameter for each trial as the distance between a bin of pre-movement neural activity and farthest point of the neural trajectory during movement (Methods). An example trial from Monkey E is shown. (D) Greater rewards generally led to greater trajectory diameter. We calculated median (± SE) trajectory diameter as a function of reward and compared values for Small and Large trials (Wilcoxson rank sum test, *** = p < 0.001, n.s. = not significant). (E) Trajectory diameter correlates with vigor (n = 489 day-direction-reward conditions). We show a histogram of the within day-direction-reward condition correlations between single trial trajectory diameter and peak speed (median r = 0.19). The bar at the top indicates the 99% confidence interval of the distribution’s median. (F) We calculated peak normalized neural speed for each trial by normalizing the neural speed in the principal components space by the trajectory diameter each trial (see Methods for details). (G) Peak normalized neural speeds increase as a function of reward. Same format as panel D. (H) Peak normalized neural speed correlates with vigor. Same format as panel E (n = 489, median r = 0.10).

To quantify the magnitude and speed of the neural trajectory, we first calculated the single-trial “trajectory diameter” as the Euclidean distance between pre-movement neural activity and the farthest point of the neural trajectory during the movement in the 2D plane (**Fig. 6C**, Methods). We observed significant increases in trajectory diameter as a function of reward in two out of three animals (**Fig. 6D**). We also found a robust trial-to-trial correlation between trajectory diameter and peak speed across all animals (**Fig. 6E**).

We next calculated the speed at which the neural trajectory was traversed on a given trial relative its trajectory diameter (**Fig. 6F**, Methods). We found that this quantity was greater for higher magnitude rewards (**Fig. 6G**) and significantly (albeit weakly) correlated with trial-by-trial peak speed (**Fig. 6H**). In summary, we found both the magnitude and speed of the neural trajectory during movement were significantly modulated by reward and were correlated with movement speed, providing another avenue through which reward may modulate motor cortex to drive movement vigor.

### Links between motor cortical correlates of reward-mediated vigor

Does reward drive vigor through one motor cortical mechanism, or multiple? We have identified seven different single-trial neural metrics across the epochs of the reaching task that are related to movement vigor (**Fig. 7A**). The fact that we see reward affecting all of these seemingly distinct aspects of motor cortex activity supports the multiple mechanisms argument. However, it is also possible that these metrics are all controlled by a single latent variable related to vigor, in which case reward may be “turning one knob” to influence vigor of behavior.

**Figure 7.**
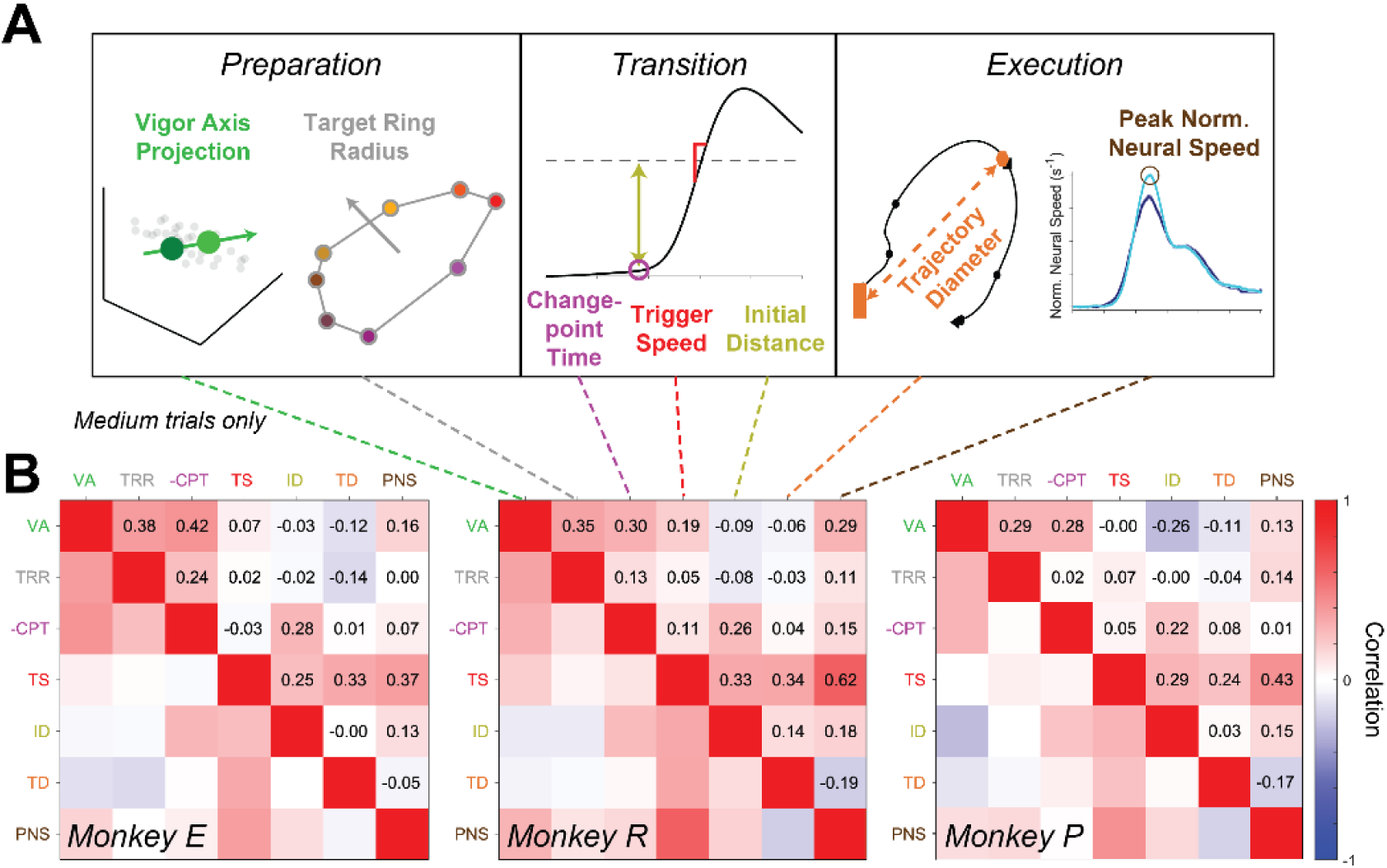
Neural metrics of vigor exhibit limited correlations with one another. (A) Summary cartoons of the seven neural metrics related to vigor identified in prior figures. (B) Spearman rank correlation heatmaps between the seven neural metrics for Medium reward trials (Methods). Note the sign of changepoint time is flipped (shown as “-CPT”) to match the convention that higher values correspond to greater vigor.

To discern between these possibilities, we calculated the correlation between each of the neural metrics that we identified. If a single latent variable could explain all the neural metrics, we would expect to see strong correlation structure; e.g., trials with higher vigor-axis projections would have a similarly greater target ring radius, faster changepoint time, etc. Contradicting this idea, we observed few strong correlations overall (**Fig. 7B**).

Even though there were not many strong correlations, we did see broad consistency in trends across animals. Three particularly salient examples are: (1) The preparatory epoch metrics and changepoint time were fairly correlated with one another (average r = 0.27), (2) trigger speed correlated with both trajectory diameter and peak normalized neural speed (r = 0.39) even though those two metrics were not positively correlated with one another (r = -0.14), and (3) initial distance, where higher values indicate neural preparatory states farther from trigger axis movement threshold, correlated with both changepoint time and trigger speed (r = 0.27). Nevertheless, the lack of strong correlation structure between the neural metrics supports the argument that reward affects multiple distinct aspects of motor cortical activity to invigorate movements.

### Reward affects more in motor cortex than correlates of vigor

We have demonstrated that reward influences motor cortex activity. But for what purpose are these reward signals present? One possibility is that they solely serve to invigorate movements. In other words, motor cortex would not actually encode reward, it would only encode vigor, which correlates with reward. A strong test of this hypothesis would be if the neural signals of vigor fully accounted for reward’s influence on motor cortex. We assessed this by subsampling trials to create vigor-matched reward conditions (**Fig. 8A**; Methods) and quantifying whether we could still decode reward from neural activity. If vigor accounts for a substantial portion of the reward signals present in motor cortex, then matching the vigor distributions between two different reward conditions should severely curtail the ability to decode reward identity from neural activity.

**Figure 8.**
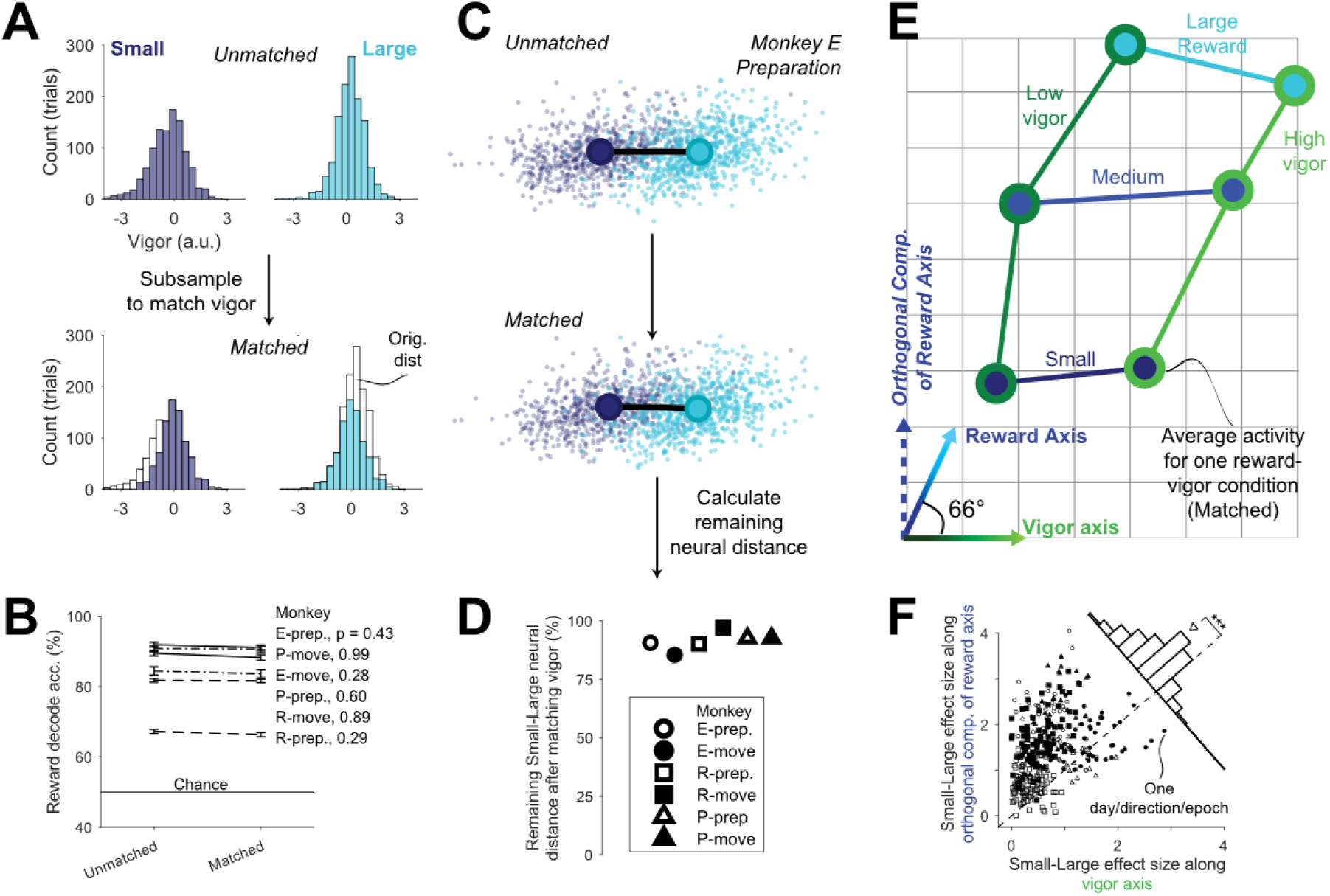
Vigor does not account for most of reward’s effect on motor cortical activity. (A) To control for the effects of vigor, we repeatedly subsampled trials from each reward condition to match vigor distributions (see Methods for details). This produced vigor “matched” subsamples, which we compared against “unmatched” subsamples that were drawn from the original vigor distributions. (B) Controlling for vigor has little impact on reward information in motor cortex. We show the average cross-validated LDA decoding accuracy (± S.E. over folds) of Small versus Large reward labels for the vigor unmatched and matched subsamples for both the preparatory epoch and the movement epoch (Methods). We see no statistically significant difference between unmatched and matched samples (p-values on plot are Welch’s t-test comparisons; see Methods). (C) We further assessed how controlling for vigor impacted neural representations of reward by evaluating how much of the Small-to-Large distance in neural activity space remained after matching vigor distributions. To visualize this, we show Monkey E’s preparatory neural activity from vigor unmatched (top) and matched (bottom) subsamples in a 2D plane that captures reward-related separation (Methods), where each small dot is one trial colored by reward (Small is dark blue, Large is light blue). Qualitatively, we see little decrease in the distance between the Small and Large trial averages after matching vigor. (D) The majority of the neural distance between Small and Large rewards remains after matching vigor distributions. For each animal and each epoch, we calculated the neural distance between Small and Large reward trial averages in both the unmatched and matched subsamples, and calculated what percentage of the unmatched value remained in the matched subsample (Methods). (E) Reward encoding in motor cortex has a large orthogonal component to vigor encoding. We calculated a “reward axis” in a similar fashion to the vigor axis described in Figure 2 (Methods), then compared the two axes. Points show the average projections for each reward-vigor condition from matched subsamples in a 2D plane that captures both the reward and vigor axis, where the inner color indicates reward label (blues) and the outer color indicates vigor label (greens). We connect points from the same vigor or reward condition for visualization. We also show the vigor and reward axes at the bottom-left, including the angle between them. The y-axis in this plot is thus the component of the reward axis that is orthogonal to the vigor axis. Quantification of the angles between reward and vigor axes are shown in **Figure S6**. (F) The effect size of reward along the vigor axis is smaller than that along the orthogonal component of the reward axis. We calculated Small to Large reward effect sizes (Cohen’s d) in the projections along both axes within each day-direction-epoch condition, then plotted them against each other (same legend convention as panel D). The inset histogram shows the difference between the two, where 280 out of the 324 conditions (86%) show greater effect size along the orthogonal component of the reward axis (median = 0.68, sign-rank test p < 10^-10^).

We found that reward decodability was not significantly different between vigor-matched and vigor-unmatched subsamples (**Fig. 8B**). To further build this intuition, we also calculated the neural distance between Small and Large reward trial averages in both subsamples. If vigor accounted for all reward effects, we would expect that the act of matching vigor would cause the two distributions to overlap, removing most of the distance between the means. Instead, we observe that the overwhelming majority of neural distance between Small and Large averages remains after vigor matching, ranging from 85% to 97% of the distance when vigor is not accounted for (**Fig. 8C-D**). Our analyses clearly indicate that vigor alone cannot account for the majority of reward information present in motor cortex.

To gain a fuller picture of the arrangement of vigor and reward signals in neural population activity space, we identified both the vigor axis and a reward axis from preparation and execution epochs of neural activity separately (Methods). The reward axis was found using the difference between vigor-matched Small and Large reward trials, and the vigor axis was again identified as in Figure 2 above, using trials of Medium reward conditions only. We found the reward and vigor axes were neither completely orthogonal nor co-linear, forming an acute angle with one another (**Fig. 8E**; see **Fig. S6** for quantification). Importantly, this means that there is a substantial portion of reward-related variance in motor cortical population activity that lies orthogonal to vigor-encoding patterns.

To get a sense of the strength of reward encoding in the component that is orthogonal to vigor, we compared the effect size of reward-driven difference in neural activity along the component orthogonal to vigor with the effect size along the vigor axis. Across animals and epochs, the effect of reward was significantly weaker along the vigor axis than along the orthogonal component of the reward axis (**Fig. 8F**). This means reward’s effects on motor cortex are more substantial in dimensions *not* related to movement vigor. Our results support the conclusion that vigor is unable to explain the majority of reward signals in motor cortex.

## Discussion

Reward invigorates movements^1,12,22^, and neural correlates of both reward and vigor have been identified in motor cortex^30,35,36,44,47,53^. Here, we examined those neural signals together. We found that reward invigorates movement through multiple neural processes. During movement preparation, we found both target direction-specific and target direction-independent manifestations of vigor. During movement initiation, we found that trials with greater vigor had shorter neural latency to the go cue and faster neural transition dynamics. Lastly, during the execution of the reach, we found that more vigorous movements both stretched neural trajectories and sped up the movement along them. These neural correlates of vigor were only loosely correlated with one another, which suggests that reward invigorates movement through multiple neural mechanisms. Finally, we observed that the influence that reward has on motor cortical activity is quite large: Even when we account for reward-mediated vigor, we cannot explain the bulk of reward’s influence on motor cortex activity.

The observation that a great deal of reward-related modulation in motor cortex was not related to vigor (**Fig. 8**) is corroborated by prior work demonstrating similar reward modulation even in animals that are not moving their limbs^14,46,47,53^. What might be the purpose of these non-vigor-related reward signals in motor cortex? One possibility is that, in this task, reward may be affecting other aspects of behavior through motor cortex that are not visible in movement vigor, such as accuracy or susceptibility to false starting^12,20^. It could also be that reward information has other roles in motor cortical activity beyond affecting behavior immediately. This would pose the excess reward information in motor cortex as a contextual signal, potentially useful for engaging neural dynamics specific to the situations indicated by the contextual reward cues^43,46,63,69–72^. In a similar vein, a third possibility is that this signal may be related to dopamine projections and potentially serve some reinforcement learning role^52,53,73^. This idea is supported by the both direct and indirect connections that exist between the brain regions that produce dopamine and motor cortex^74–79^.

Reward-mediated vigor is ubiquitous across many tasks and species^1,3,11,12,18,35^. There are arguably two distinct and complementary reasons why greater reward often increases movement vigor. The first reason is *innate*. For even the simplest lifeforms grabbing nutrients in an uncertain and dangerous environment, avoiding the penalty of loss (or death) encourages extra expenditure of effort to get bigger rewards more quickly. One could then argue that this notion of reward invigorating movements is baked into the evolutionary development of the nervous system, tethering together the value and motivation behind an action with its vigor^80–82^. The neuromodulator dopamine provides supporting evidence for this innate argument of reward-mediated vigor, as it serves jointly important roles in value encoding and driving movements^25,66–73^. Multiple studies using saccades^2,9,10^ and reaching^20^ have demonstrated movement vigor modulated by reward even when the task at hand was not benefited by it, in line with an innate explanation of reward-mediated vigor.

The second reason reward may invigorate behaviors is *strategic*. Manohar and colleagues^6^ proposed that greater reward effectively allows more effort to be expended for a given action - that is, greater reward pays the cost of increased effort. A strategic argument would then be that these greater rewards permit an individual to allocate more effort into some aspect of volitional control, and in this task, vigor is a strategic quantity to choose. This idea could also account for experimental evidence of how reward can also improve accuracy alongside vigor improvements^6,12^ along with potentially explaining inter-subject differences in reward-driven behavioral changes.

In our experiment, both the innate and strategic drivers of reward-mediated vigor operate in the same direction: move faster for greater rewards. But, consider tasks where the two are *not* aligned. These could be tasks where increased vigor is a secondary objective to performance (e.g., typing), tasks where vigor is neither helpful nor harmful (e.g., free throws in basketball), or tasks where increases in vigor could be detrimental (e.g., musical performance). In many cases, we would want reward to improve movement precision, an aspect often at odds with vigor^74^. This push/pull of reward-driven vigor and precision may account for the “choking under pressure” we observed in prior work, where we found that when animals were presented with rare and gigantic rewards, they would reach with excessive caution (low vigor) and fail more frequently^35,83^.

Considering the interplay of innate and strategic drivers of reward-mediated vigor summon many questions that both directly and indirectly build off the work presented here: Do aspects of reward encoding in motor cortex flexibly adapt to task requirements, or are they static in their relationship with movement vigor? How do we allow the “innate” sense to invigorate us when we need to push past our limits, yet suppress it so we can perform our best in other highly motivating cases? Are all creatures capable of this flexible strategic-versus-innate modulation of behavior with reward, or does it require certain cognitive and neural faculties? We view our study as a stepping stone to probing these questions and more to unveil the neural mechanisms underlying how motivation controls behavior.

## Methods

### Subjects

Three adult male rhesus macaques, Monkeys E (9.0 kg, 10 years old), P (9.5 kg, 6 years old), and R (19.0 kg, 10 years old) performed the experiments in this work. All experimental and animal procedures were approved by the University of Pittsburgh Institutional Animal Care and Use Committee in accordance with the guidelines of the US Department of Agriculture, the International Association for the Assessment and Accreditation of Laboratory Animal Care, and the National Institutes of Health.

### Task

The experimental setup used in this study is described in full in Smoulder et al. 2024^35^ (**Fig. 1A**). Briefly, the three monkeys in this study were on standard water regulation procedures to maintain motivation and valuation of liquid rewards. During experiments, each monkey sat in a primate chair facing a mirror approximately 8 centimeters from his eyes that reflected a computer screen displaying task events. The monkeys performed tasks by moving their hands in the open space in front of them behind and below the mirror (Monkey E and R right hand, Monkey P left hand). We tracked hand position in 3D using an infrared LED marker attached to either the monkey’s working hand’s index finger (Monkeys E and P, 120 Hz sampling rate, nominal resolution < 1mm) or back of the hand (Monkey R, 60 Hz sampling rate, nominal resolution < 1mm). The monkey’s hand position in the frontoparallel (coronal) plane corresponded 1-to-1 with cursor position on the screen. Any trials where tracking of the hand failed at any point in time were removed (< 1% of trials).

The monkeys performed a delayed center out reaching task, described in detail in Smoulder et al. 2021^83^ (referred to in that work as the “speed + accuracy task”). Specific task parameters for the subjects in this work can be found in Smoulder et al. 2024^35^, though we re-list particularly relevant parameter values as they are described here. A trial began with a circular target appearing at the center of the display. The animal initiated a trial by moving the cursor into this center target and holding for 200-600 ms. After this, a reach target appeared at one of multiple possible locations a set distance away from the center (Monkey E: 85 mm, R: 80 mm, P: 65 mm). For Monkeys E and R, these were 8 equal-angular spaced targets in a circle around the center (i.e., right, up-right, up…). For Monkey P, these were 4 equal-angularly spaced targets in a square (up-right, up-left, down-left, down-right). The reach target was visible for the remainder of the trial.

Either the color (Monkeys E and P) or the image (Monkey R) inside of the reach target cued the animal as to the reward size at stake for the trial. Reward values for Small, Medium, Large in mL were: Monkey E: 0, 0.2, 0.4. Monkey R: 0, 0.22, 0.44. Monkey P: 0.15, 0.3, 0.6. The animals demonstrated understanding of the relative value of the cues in a separate two-target choice task; see Smoulder et al. 2024 for details of animal training, reward cue learning, and choice behavior^35^.

After the reach target with the reward cue appeared, a variable-length delay period began (ranging from 200-950 ms across animals) during which the animal had to keep the cursor within the center target. When the delay period elapsed, the center target disappeared as a “go cue” for the animal. The animal then had a short period of time to move the cursor to land in the reach target (Monkey E: 725 ms, R: 625 ms, P: 850 ms). Once they moved the cursor into the reach target, they had to hold it inside for 400 ms, after which they received the cued reward. Unsuccessful trials could be due to delay failures (defined as the cursor exiting center target before 100 ms after the go cue), reach failures (which occurred when the cursor either did not make it to the target in the allotted time or blew straight through it) or target hold failures (failing to hold in the reach target for the full 400 ms after the hand stopped inside for at least 150 ms). If the animal failed a trial, the cursor and targets turned purple and the screen froze for at least 1 second, after which the animal had to make an unrewarded reach to another random target (this reach had no time limits). This unrewarded reach discouraged the animals from quitting Small reward trials. After this, the animal could proceed with the next trial.

### Data collection and preprocessing

#### Behavioral data and vigor computation

The kinematic data that we analyze in this work are based on the cursor position signals that the animal controlled with their hand during the task. We smoothed cursor position signals with a zero-phase low pass filter (2nd order with 10 Hz cutoff), then calculated cursor velocity from the raw data using the time derivative of the cursor position in the 2D task plane. For each trial, we then linearly interpolated the cursor position and velocity signals up to 1000 Hz, assuming each raw cursor velocity sample was timestamped at the midpoint between the two position values used for its calculation. Cursor speed was then calculated as the square root of the sum of squared horizontal and vertical cursor velocity (**Fig. 1B**).

Using cursor speed, we calculated two key metrics related to movement vigor for each trial (**Fig. S1A**). First, we calculated *peak speed* as the maximum cursor speed occurring after go cue and before either target acquisition. To calculate *reaction time*, in each trial we identified the earliest time point where the cursor speed was above 45 mm/s, working backwards from the time of peak speed. We chose this as a static threshold to use because (1) it does not tether the reaction time of a trial to its peak speed as it would if the threshold were determined as a percentage of peak speed, helping to decouple the two values, (2) using a static threshold permits intuitive comparison across animals and conditions, (3) it is similar to threshold values used in prior work^35,83^, and (4) it is a close match to intuitive percentiles of the animals average successful trial peak speed (Monkey E: 10% of average successful trial peak speed = 47mm/s, Monkey R: 10% = 44 mm/s, Monkey P: 20% = 46 mm/s). We also tested the analyses in this work using other thresholds (e.g., 10% or 20% of average peak speed per animal) and observed no meaningful differences in results. Once we identified the earliest time point of cursor speed being above threshold, we linearly interpolated time between it and the preceding index to identify the time value that corresponded to 45 mm/s. This was then saved as the reaction time for that trial. We generally found that reaction time and peak speed were weakly correlated with one another (**Fig. S1B-C**).

Because prior work has demonstrated improvements in “vigor” by both the speeding up of reaction times and peak speed, we define vigor for this work as the combination of the two as shown in equation [1]. We z-scored values of peak speed and (inverted) reaction time within each day-direction condition, agnostic to reward. (Note two exceptions to this: First, in **Figure 1D** we use z-scored values across all trials for each animal to show variability across days, and second, when reach direction neural signals were being compared, as described below in *Target plane analysis*). We invert reaction time to give it the same sign as peak speed (i.e., higher value corresponds to greater vigor), and because inverse reaction times had a more normal distribution in our data than reaction times did^59^. We studied kinematic metrics preceding and up to the time of peak speed to focus on the ballistic portion of the reach, as the targets in this task were small and necessitated careful approach after this ballistic launch^83^.

#### Neural data

All animals in this study were implanted with multielectrode “Utah” arrays (Blackrock Microsystems, Inc). We recorded neural activity from the primary motor cortex (M1) and dorsal aspect of the premotor cortex (PMd) using implanted multielectrode arrays. Monkey E had one 96-electrode array that straddled the shoulder regions of M1 (∼64 channels) and PMd (∼32 channels). Monkey R’s recordings were from a 96-channel array in the shoulder region of M1. Monkey P’s recordings were from two arrays, 32 channels in M1 and 64 channels in PMd.

Because recordings in both regions were responsive during this task and showed fairly heterogeneous tuning to task parameters, we make no distinction between the two, referring to the combined M1 and PMd recordings as “motor cortex.” All recordings were bandpass filtered from 300 to 5000 Hz to isolate spiking activity.

Before each experiment began, we set voltage thresholds on each electrode at a multiplier of the root-mean-square voltage (Monkey E: -3.5x, P: -3x, R: -3x)^84^. We stored 1 ms waveform snippets (sampling rate 30 KHz) surrounding each threshold crossing during the experiment. Individual units were manually sorted on each electrode using offline spike sorting (Plexon). Units were identified visually using a combination of features, including waveform principal components and peak-to-trough amplitude. We included both well-isolated units and multi-unit waveforms for this study. We excluded units with mean firing rates of less than 1 spike per second.

### Data selection and statistics

The trials of the delayed center out task analyzed in this work are a subset of those analyzed in Smoulder et al. 2024^35^. Here, we excluded four groups of trials. First, we exclude delay epoch failure trials, as they do not have accurate measures of reaction time and peak movement speed. Second, we exclude the rare and high-magnitude “Jackpot” reward used for 5% of trials, as the animals performed paradoxically worse for Jackpots and we believe this “choking under pressure” is a potentially distinct phenomenon from reward-mediated invigoration of movement^35,83^. Third, we exclude trials with uncharacteristically high or low reaction time and/or peak speed, which we define as exceeding 6 median absolute deviations away from the median. This was less than 1% of trials for each animal. Fourth, we exclude the first 4 sessions collected with Monkey P, as this was immediately after array implantation and consequently those sessions had more substantial recording instabilities. Including these sessions did not impact the results or interpretations of the present study.

All statistical tests performed in this work are two-tailed. Detailed statistical information for all main figures is listed in **Table S1**. In general, we performed statistical comparisons using the means of data that were qualitatively normally distributed; for those that were not, we used the median. For standard error bars, when the mean was used, we calculated standard error of the mean as the standard deviation divided by the square root of the number of observations. When the median was used, we calculated the standard error by creating a bootstrapping sample (i.e., fully resampling with replacement), estimating the median for the bootstrap sample, and then taking the standard deviation across 10000 separate bootstrap samples. For comparisons of means across groups, we use Welch’s t-test for comparing groups of unequal variance to assess statistical significance. For comparisons of medians, we use a Wilcoxson rank-sum test. We only assess statistical differences between Small and Large reward trials when assessing the effect of reward (i.e., discounting Medium reward trials), as the majority of the trends observed in this work were monotonic as a function of reward. All correlations reported are calculated using the Spearman rank correlation for robustness, though we observed nearly the same results using Pearson correlation.

### Behavioral analyses

We assessed the animals’ success rates and failure modes as a function of cued reward (**Fig. S1F**). We gave each failed trial one of five possible labels, as in prior work^35,83^ (see Smoulder et al. 2021 for failure mode details). Delay failures were classified as either false starts if the cursor speed exceeded 100 mm/s along the vector pointing from the center target to the reach target, or a delay drift failure if otherwise. Undershoot trials were classified as those where the cursor did not enter the reach target within the allotted reach period time and the cursor never passed the midline of the reach target (i.e., it did not blow past the target off at an angle). Overshoot trials were those where the cursor either blew past or through the target without stopping. Target hold failures were those where the cursor was deemed to have stopped in the reach target (speed below 50 mm/s and in target for more than 100 ms) but then exited before the required time. Outside of **Fig. S1F**, we excluded delay failure trials or trials with aberrant reaction time or peak speed from further analyses as described above in *Data selection and statistics*. Results from this study did not meaningfully change if only successful trials were used.

We assessed reaction time and peak speed as a function of reward. Because reaction time distributions often have heavy tails and are not normally distributed^59,60^, we use the median and a standard error of the median (estimated via bootstrapping, see *Statistics*) instead of mean and standard error of the mean. We pooled data across all sessions and reach-direction conditions within each animal, then calculated the median reaction time and mean peak speed along with standard errors within each reward condition (**Fig. 1C**). We also looked at the variability in these metrics as a function of reward (**Fig. S1D-E**). We then assessed how robustly reward affected vigor by calculating the average vigor within each reach-direction condition for each day and reward size (referred to as a day-direction-reward condition), then plotting the Small and Large averages against one another (**Fig. 1D**).

### Neural analyses

#### Comparing single unit reward and vigor tuning

We constructed peri-stimulus time histograms (PSTHs) for various units recorded from Monkey E for visualization. To do this, we aligned neural data within each trial to either target onset or go cue, then smoothed spike trains with a 25 ms standard deviation Gaussian kernel. To evaluate the effects of reward, we calculated the mean firing rate (and S.E.) across trials in Small and Large conditions (**Fig. 2A**). To evaluate the effects of vigor, we took the Medium reward trials and gave each a vigor label of “Low” or “High” based on whether its vigor value was below or above the median. By using only Medium reward trials in assessing vigor effects, we use a disjoint set of trials from those used in assessing reward effects. We then calculated mean firing rate (and S.E.) across Low and High vigor trials (**Fig. 2B**). We also more thoroughly examined single unit activity at the end of the delay epoch to assess if tuning to reward and vigor was correlated and if vigor tuning was consistent across reward conditions (**Fig. S2**).

#### Neural data stitching

We stitched neural recordings across days using a modified version of factor analysis as described in prior work^35^ and summarized in the following paragraphs. We stitched the data primarily for the statistical power of being able to combine data across days, which enables robust study of the relatively subtle effects of reward and vigor on motor cortical activity. As with any other factor analysis model, the stitched neural data is a linear projection of the single units firing rates. The stitched neural data was used for all analyses aside from those designating single units (**Fig. 2A-B** and **Fig. S2**).

At a high-level, neural stitching involves first identifying units that were chronically recorded across consecutive days, then performing factor analysis with missing entries filling in for days when a given unit was not present. For identifying units that were present across multiple sessions, we implemented the method from Fraser & Schwartz 2012 that uses mean firing rate, waveform shape, spike time autocorrelation, and spike time cross-correlation to predict if a unit on a given channel is the same across days^85^. We used a false positive threshold of 0.99 (i.e., for a unit to be called “same” across sessions, it has to be outside the 99% percentile of a distribution of similarity statistics calculated from different channels), which errs on the side of caution in calling units the same across days. We then binned spikes from each trial in non-overlapping 200 ms bins aligned to movement onset, going as far back as 400 ms before target onset and as far forward as 400 ms after target acquisition (on average, 8 bins per trial). We mean-centered the firing rates for each unit each day by subtracting the average across all of these bins. We constructed a matrix of trial-concatentated firing rates across all sessions, where each row was a time bin and each column was a unit, and filled in blank values on sessions when a given unit was not present. We then performed factor analysis with missing values, as described in ^65^ and ^35^. We used methods described in prior work to initialize this factor analysis model to avoid local minima issues (fitting factor analysis models within each session and aligning them with Procrustes method; see ^64,66^ for details) and orthonormalized the output fitted model using singular value decomposition^86^.

The end result of this processing is the identification of a single factor analysis model that applied across all data from all sessions for a given animal. This means that the factor scores for any one session are directly comparable to the factor scores from data collected in any other session. After fitting the final model, we projected the spike rates at 1 ms resolution along the orthogonal factors. We used 10-fold cross-validation to select the number of factors to use (Monkey E: 24 factors, R: 18 factors, P: 16 factors). For neural population analyses (the remainder of the Methods), we used the factors found via the neural stitching procedure described above.

#### Calculating vigor axes in preparatory activity

We sought to identify a dimension in neural preparatory activity that captured information about upcoming movement vigor across all reach direction conditions (**Fig. 2C**). To identify this, we averaged neural activity over a time bin ranging from 150 ms before the go cue to 50 ms after. Because we wanted to analyze the preparatory “steady-state” after target onset transient neural activity settled, we excluded trials with a delay epoch of less than 400 ms for these end-of-delay analyses only. We again used only data from Medium reward trials, though instead of z-scoring vigor and neural activity by day-direction, we only mean-centered. This is because z-scoring neural activity in the form of factor scores would improperly assign each factor equal variance, and we wanted to perform the same operation on vigor and neural activity. We then again split these trials into Low and High vigor based off of their median, and calculated the average neural activity for these Medium reward Low and High vigor trials. We defined the vector that connected the Low and High vigor means as the “vigor axis” and projected all trials’ neural preparatory activity along it to produce *vigor-axis projections*.

For all neural metrics identified in this work, starting with the vigor-axis projections, we performed the same procedure to assess modulation as a function of reward and vigor. To assess the effects of reward, we calculated the average value as a function of reward and statistically compared Small and Large reward averages (**Fig. 2D**). To assess the effect of vigor, we calculated the correlation of the neural metric with the behavioral measure of vigor in each day-direction-reward condition that had at least n = 10 trials. We then made a histogram with these correlation values and calculated their median along with a 99% confidence interval of the median. Because Medium reward trials were directly used to identify vigor-axis projections and this “bakes in” the correlation between neural activity and behavior on those trials, we exclude them from this analysis specifically (**Fig. 2E**). The combination of these two analyses demonstrates that a given neural metric is both modulated by reward and correlates with movement vigor across different task conditions. Along with the method described above, we tested other ways of identifying the vigor axis and compared them to justify using this method in the main text (**Fig. S3**), though all produced comparable results.

#### Calculating target ring radius from preparatory activity

To study how reward and vigor relate to direction-specific preparation signals in motor cortex, we first used LDA to identify a 2D plane of neural activity that best separated the cued target directions (**Fig. 3A-B**). We visualized activity in this plane by taking the average of the neural activity as a function of reach direction and either vigor label (**Fig. 3D**, medium trials only) or reward label (**Fig. 3C**). To quantify the separability of reach direction conditions in the target plane projections, we calculated the *target ring radius* (**Fig. 3E**). This metric is equivalent to the “target preparation axis projection” of prior work^35^. To find this, we first calculated the average activity for each direction condition in the target plane (intuitively, the points on the ring). We also calculated the average of these direction averages (the center of the ring). We then took neural activity for each of the trials for a given reach direction and projected it along the vector connecting the center of the ring to its average for that direction, effectively quantifying a single-trial metric of separability from the center. We then z-scored values within each day-direction condition before combining them. We calculated average values as a function of reward (**Fig. 3F**) and also calculated correlations with vigor within each day-direction-reward condition (**Fig. 3G**). Results did not meaningfully change if the target average rings were calculated within reward/vigor condition instead of across all trials.

#### Trigger axis and transition epoch analyses

We wanted to examine how reward and vigor related to movement initiation, occurring between movement preparation and execution. To study this epoch, we focused on a “trigger axis”, a dimension of motor cortical activity that shows a reliable and robust condition-invariant signal whose dynamics predict reaction time (**Fig. 4A-B**)^36,38,40,68^. To begin, we first smoothed the neural activity with a 35 ms standard deviation Gaussian kernel. We use a wider kernel here than before to accommodate the noise incurred in taking the time derivative in later analyses. With these smoothed trials, we then proceeded to find the trigger axis as follows: First, we identify the optimal neural data time bins for identifying the trigger axis. Next, we establish the trigger threshold value. Finally, we quantify single trial metrics that determine when the neural signals hit the trigger threshold.

To identify the optimal neural data time bins, we start with a 50 ms bin of neural activity well-preceding movement (“bin 1”) and a 50 ms bin near the time of movement onset (“bin 2”) on each trial, balancing trials by direction-reward condition. We then performed LDA to find a dimension of maximal linear separation between bin 1 and bin 2. This dimension is a putative trigger axis. We projected all trials’ smoothed neural activity ranging from 100 ms before the go cue to 600 ms after along the putative trigger axis. We then swept a threshold criterion across values of this axis. For each threshold value, we evaluated when each trial’s neural signal first exceeded the threshold, enforcing a minimum value of 50 ms after the go cue and a maximum of 450 ms. We also noted the fraction of trials that did not cross the threshold in this interval. Finally, with all trials that did cross the threshold, we calculated the correlation of the time of threshold crossing (“trigger time”) with reaction time.

We performed a grid search over bin 1 values, bin 2 values, and threshold values and calculated the fraction of trials that crossed threshold and the correlation of trigger times with reaction time. We swept values for the center of bin 1 ranging from 400 to 0 ms before movement onset in 25 ms increments, bin 2 ranging from 200 ms before to 200 ms after movement onset in 25 ms increments (with a minimum of 100 ms between bins), and threshold along 100 evenly spaced values from 20% to 80% of the range of the average projection along each putative trigger axis. We selected the final combination of these three parameter values that maximized the quantity *rf^2^*, where *r* is the correlation value of trigger times and reaction times, and *f* is the fraction of trials that validly crossed the threshold. We square *f* to strongly penalize excluding trials, as many threshold values show similar correlation values. In general, selected threshold values ended up being near 40% of the range. For Monkey E, final bin values were (with respect to movement onset) [-375, -325] ms for bin 1 and [50, 100] ms for bin 2, with 33 trials excluded (0.91%). For Monkey R, [-400, -350] ms and [25, 75] ms with 5 trials excluded (0.05%). For Monkey P, [-350, -300] ms and [-125, -75] ms with 13 trials excluded (0.70%). Final correlation values of trigger time and reaction time for non-excluded trials are shown in **Figure 4C**. To assess the effects of reward on trigger time, we calculated the median value within each reward condition and compared Small and Large reward medians (**Fig. 4D**). We use median instead of mean for the same reasons as reaction time.

Having identified a trigger axis for each animal, we next considered multiple aspects of the trigger signal on each trial that could contribute to it hitting the threshold more quickly. We calculated three quantities for each trial (**Fig. 5A**). First, we calculated the *trigger speed*. We took the time derivative of the trigger axis projection on each trial and calculated its maximum value on an interval of 75 ms before to 75 ms after trigger time, as this is approximately when peak trigger speed occurred generally. We then z-scored values within each day-direction condition. Second, we calculated the *changepoint time* by evaluating when the trigger speed on a given trial exceeded 30% of the median peak trigger speed across trials. Like evaluating reaction time with hand cursor speed, we worked backwards from the time of peak trigger speed to identify the changepoint time. We set a minimum value of 25 ms after go cue, which excluded a handful of outlier trials (Monkey E: 2.0% of trials, R: 0.4%, P: 3.8%); including these trials did not meaningfully affect results. We did not z-score changepoint time to maintain interpretation of its units (ms), though subtracted the median value for plotting to facilitate showing all animals’ data simultaneously (Monkey E: 87.5 ms, R: 151.5 ms, P: 99.8 ms). Third, we calculated the *initial distance* from threshold as the difference between the trigger threshold and the trigger axis projection value 150 ms before the changepoint time; higher values indicate greater distance from threshold. We then z-scored values within each day-direction condition. For visualization, we show Monkey E’s average trigger axis projections over time (**Fig. 5B**).

For each metric, we calculated the average across all trials within each reward condition (**Fig. 5C,E,G**; median for changepoint time, mean for others) and compared values between Small and Large rewards. We also calculated the correlation with vigor within each day-direction-reward condition that had at least 10 trials available (**Fig. 5D,F,H**). We note that all significant trends listed in these figures are also significant within each individual animal (see **Table S1**). We further note that trigger axes have been identified in multiple different ways in prior work, and we tested three alternative approaches here as well that produced similar results (**Fig. S4**).

#### Movement epoch analysis

We analyzed how reward and vigor affected motor cortical activity during movement execution. Because we align data to movement onset for movement epoch analyses, which is after the reaction time, we use the z-scored peak speed as our metric of vigor for the movement epoch. Using the combined vigor metric shows similar, albeit weaker, results. Motor cortex activity during movement is highly dynamic, unlike the steady-state achieved during preparation. As such, we studied aspects of the neural trajectory that unfolded over the time immediately preceding movement onset through the end of the arm movement. Based on prior work, we sought to assess how reward and vigor related to two key aspects of the neural trajectory during movement: the magnitude of the trajectory and the relative speed at which the neural trajectory was traversed. Without adopting a strict rotational dynamics framework (e.g., Churchland et al. 2012^44^), this can be intuitively thought of as being similar to the amplitude and rotational velocity of the neural trajectory.

To calculate these quantities, we first smoothed neural activity using a 35 ms standard deviation Gaussian kernel. We then identified dimensions of neural activity that captured temporal variance by performing PCA on the neural trajectory (averaged across all trials) from 400 ms before movement onset to 400 ms after. We projected all trials onto the top two principal components (**Fig. 6A-B**), as these captured over 90% of the temporal variance in the average trajectory; we note that including all dimensions instead of just the top two PCs did not meaningfully affect results. For each trial, we then calculated the *trajectory diameter* as the maximum Euclidean distance in the 2D PC space between pre-movement neural activity (averaged for each trial from 400 to 300 ms before movement onset) and any point in the rest of the trajectory (**Fig. 6C**). The time of maximum distance typically occurred near or shortly after movement onset time. Like previous metrics, we compared values for Small and Large reward trials (**Fig. 6D**) and calculated within day-direction-reward condition correlation with peak speed (**Fig. 6E**).

We also assessed the relative speed of the neural activity traversing the trajectory on each trial. We considered multiple methods to assess this (e.g., KiNeT^71,87^, rotational velocity^44^), though opted to stick to a more direct metric that leverages methods used in calculating other values in the present work. We first calculated the neural speed as the magnitude of the time derivative of the neural activity in the 2D PCA space identified for the trajectory diameter above. At this stage, this is effectively a “translational” neural speed, which is affected both by the trajectory diameter and the relative speed at which the trajectory is being traversed. To isolate relative speed, we divided the neural speed by the trajectory diameter on each trial (**Fig. 6F**). Finally, we calculated the peak value of this normalized neural speed for each trial, giving us *peak normalized neural speed* values. We note that without the normalization step, peak neural speed is highly correlated with trajectory diameter (**Fig. S5A**); including it, the two exhibit weak correlation (**Fig. S5B**). We compared these values between Small and Large reward trials (**Fig. 6G**) and calculated within day-direction-reward condition correlation with peak speed (**Fig. 6H**).

#### Comparing neural metrics of vigor

We identified seven single trial neural metrics in the main text figures that are modulated by reward and exhibit putative relationships with vigor. To recap, these are (**Fig. 7A**):

1. Preparatory activity vigor-axis projections (similar to metrics used in ^30,34^)
2. Target ring radius (used in ^35^)
3. Changepoint time (similar to a metric used in ^36,40^)
4. Trigger speed (similar to a metric used in ^40^)
5. Initial distance (similar to a metric used in ^30^)
6. Trajectory diameter (similar to a metric used in ^44^)
7. Peak normalized neural speed (similar to metrics used in ^40,45^)

While all of these metrics are based on prior work that studies the relationship between motor cortical activity and some aspect of movement vigor, they have not (to our knowledge) been directly compared with one another. As such, we do not know if their relationship with vigor can be captured by a single underlying latent variable, or if there are multiple independent sources of vigor-related information available among them. To explore the possibility that one variable can explain these metrics, we examined the correlation between their values on Medium reward trials (**Fig. 7B**), expecting that if a single latent variable explained them that we’d observe strong correlation structure.

#### Evaluating reward signals after controlling for vigor

We considered the hypothesis that all reward information present in motor cortex population activity could potentially be explained by reward-mediated vigor differences. To test this, we first needed to subsample our data such that different reward conditions had the same vigor distributions. In this type of vigor-matched subsample, any remaining reward information present would thus not be attributable to a difference in vigor distributions between the reward conditions, as the vigor distributions are the same.

To create vigor-matched subsamples within each animal, we made histograms of vigor values for each reward using 20 evenly-sized bins ranging from the minimum to the maximum vigor value. To ensure proper sampling for each reward, we excluded any bins that had less than 5 trials for any reward condition (e.g., a very low vigor bin might have 10 trials for Small rewards, but only 2 for Large; this bin would be excluded). These trimmed histograms are what is shown in the top of **Figure 8A** for Monkey E. From here, for a given matched subsample, we stepped through each vigor bin and randomly subsampled a number of trials equal to the minimum count across reward conditions. For example, if Small has 18 trials for a given vigor bin, Medium has 15, and Large has 12, we would randomly choose 12 trials from each reward condition. By doing this, we produce a subsample of trials that has identical vigor histograms for each reward condition (**Fig. 8A**, bottom). We performed this operation 200 times to get many vigor-matched subsamples. Overall, this subsampling procedure excluded 29.3% of trials for Monkey E, 14.8% of trials for Monkey R, and 20.4% of trials for Monkey P. For comparison, we also performed 200 vigor-unmatched subsamples, which had an equal number of trials to the matched subsamples, but were randomly drawn from the trimmed histograms’ data.

If vigor could wholly account for reward information in motor cortex, then reward should not be decodable from the vigor-matched subsamples. To test this, we performed 10-fold cross-validated LDA decoding of reward identity (only using Small and Large trials) from the vigor-matched and vigor-unmatched subsamples. We performed this operation on the same bin of preparatory activity studied earlier from the end of the delay epoch (“Prep.” in **Fig. 8B**), along with a peri-movement bin of neural activity from 50 ms before movement onset through 150 ms after (“Move” in **Fig. 8B**). For each subsample, we calculated the mean, standard deviation, and standard error of the mean over the decoding accuracies calculated in each of the 10 test folds. We then averaged these values across subsamples to get a single value of each for the vigor-matched and vigor-unmatched subsamples within each animal-epoch. To assess if there was a statistical difference in the mean decoding accuracy between the vigor-matched and vigor-unmatched subsamples for a given animal-epoch, we performed a 2-tailed Welch’s t-test that used the average decoding accuracy means as sample means, and the standard deviations over folds as sample standard deviations, with 18 degrees of freedom (10 folds for matched subsamples, 10 folds for unmatched, minus 2).

For further intuition on how controlling for vigor impacted reward encoding in motor cortex, we evaluated the neural distance between Small and Large rewards in both the vigor-matched and vigor-unmatched subsamples. If matching vigor across reward conditions eliminated neural reward information, then we would expect the Small and Large reward averages to effectively be on top of one another (distance ∼= 0) in the vigor-matched subsample. To visualize this, we show datapoints from one unmatched and matched subsample in Monkey E’s data projected in a 2D plane, where the horizontal axis captures the full difference between reward means and the vertical axis is the orthogonal dimension that explains the most trial-to-trial variability in the data (**Fig. 8C**; the vertical axis is solely for visualization). We calculated the average of this neural distance across all subsamples, then calculated the average percentage of distance that remained in the vigor-matched subsample (**Fig. 8D**)

#### Comparing reward and vigor axes in motor cortex

After the preceding analyses studying how neural correlates of vigor were modulated by reward, we considered how reward information could coexist with vigor-related signals in motor cortex. We identified a “reward axis” in a similar manner to the vigor axis described above: using vigor-matched subsamples, we found the average vector that connected Small and Large reward average activity in the neural population activity. We did this for each animal in the same preparatory and movement bins as for decoding reward. We also calculated vigor axes using Medium reward trials for the same preparatory and movement bins. We quantified the angle between these axes within each epoch (**Fig. S6**).

For visualization, we plotted Monkey E’s average activity for each reward condition and Low versus High vigor in a 2D space spanning the preparatory reward and vigor axes (**Fig. 8E**). The vertical axis of this projection was found using QR decomposition to identify the orthogonal dimension to the vigor axis with minimal angular deviation from the reward axis. We accordingly call it the orthogonal component of the reward axis, shortened for the remainder of methods to “orth. reward axis.”

We wanted to assess how strong reward signals were along the vigor axis relative to the orth. reward axis. To do so, we projected all trials’ data from the preparatory and movement epochs along their respective vigor axis and orth. reward axis. We then divided the data into day-direction groups and calculated the effect size (Cohen’s d) between Small and Large reward trials for all groups where at least 10 trials were present (n = 324 day-direction-epoch conditions). We plotted orth. reward axis effect sizes against vigor axis effect sizes, then made a histogram of the differences and evaluated if the median significantly deviated from 0 (**Fig. 8F**).

## Data and code availability

MATLAB code used for generating figures is located at https://github.com/adam-smoulder/Reward-influences-movement-vigor-through-multiple-motor-cortical-mechanisms/. Data will be made available upon reasonable request.

## Acknowledgements

This work was supported by: National Institutes of Health grants R01NS129098 (APB,SMC), R01NS129584 (APB,SMC) and CRCNS R01NS105318 (APB), National Science Foundation grants DGE1745016 (ALS), DGE2139321 (PJM), and DRL2124066 (SMC) / DRL2123911 (APB), the Achievement Rewards for College Scientists’ Pittsburgh Chapter Award (ALS), and the Bradford and Diane Smith Graduate Fellowship (ALS).

## Author contributions

Conceptualization: ALS, SMC, APB. Methodology: ALS, PJM, APB, SMC. Software: ALS, PJM. Validation: ALS. Formal analysis: ALS. Investigation: ALS, PJM, ERO, SES. Resources: APB, SMC. Data curation: ALS, PJM. Writing (original draft): ALS, APB, SMC. Writing (review/editing): ALS, PJM, ERO, SES, APB, SMC. Visualization: ALS, APB, SMC. Supervision: APB, SMC. Project administration: ALS, PJM, APB, SMC. Funding acquisition: ALS, PJM, APB, SMC.

## Additional information

**Supplementary information** for this work is available in the Supplemental Materials PDF.

**Correspondence** and request for materials should be directed to Aaron Batista and Steven Chase.

