## Supplementary Information for "Reward influences movement vigor through multiple motor cortical mechanisms"

This PDF includes:

Figures S1-S6
Table S1

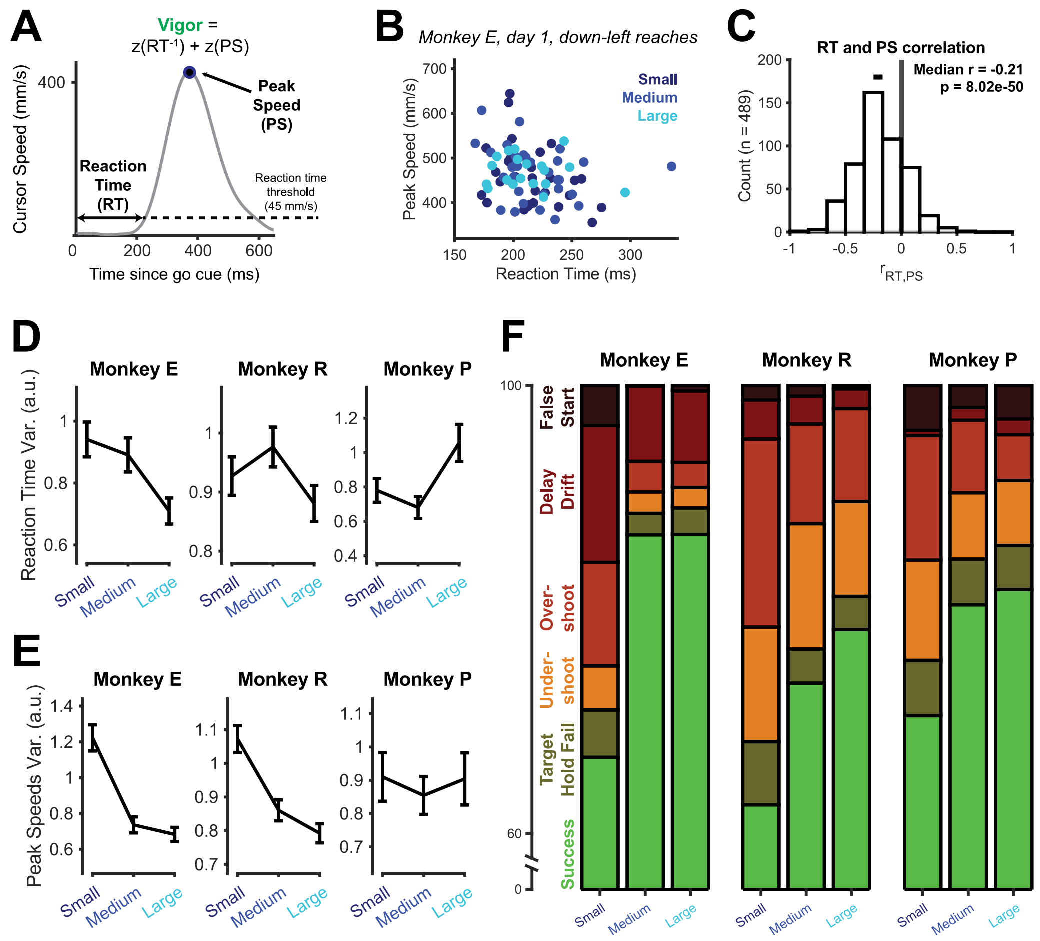


**Figure S1. Reward modulated multiple aspects of behavior related to vigor.**

To quantify movement vigor in this work, we use a normalized combination of single trial reaction time and peak speed (PS) as shown in equation [1]. Given that both are related to receiving task rewards more quickly, we wanted to explore the correlations between the two metrics.

(A) We show an example cursor speed profile from a single trial to demonstrate definitions of reaction time (RT), peak speed (PS), and vigor (where z is the z-score operation, Methods). For the reaction time, we used a static threshold of 45 mm/s across all animals (dashed line, Methods).

(B) Example scatterplot showing correlation between reaction time and peak speed for a single target on a single day.

(C) Reaction time and peak speed are significantly correlated, albeit not very strongly. We calculated the correlation between reaction time and peak speed within each day-direction-reward condition that had greater than 10 trials (n = 489). We then made a histogram of these correlation values (p-value calculated from sign-rank test). The black error bar shows a 99% confidence interval of the median across all correlation values.

We observe a small, albeit significant negative correlation between reaction time and peak speed. This is the expected direction of correlation, as faster reaction times yield smaller values. Importantly, the two are not highly correlated, meaning they both provide unique information into the quantification of vigor.

Along with looking at trends in reaction time, peak speed, and vigor as a function of reward (**Fig. 1**), we also considered that reward may affect variability in vigor metrics. To do so, we first z-scored the inverse reaction time and peak speed values with each day-direction condition. We then calculated the variance of these values for each reward. To get error bars, we performed a bootstrapping procedure: for each reward, we performed 5000 bootstraps (resampling with replacement) within each day-direction condition. We then calculated the sample variance within each bootstrap. Finally, we used the standard deviation across the 5000 bootstraps as our estimate of the standard error of the variance, which made the error bars.

(D) Reaction time variability shows subject-specific trends as a function of reward. Monkey E's reaction time appears to be less variable for Large reward trials compared to Small, while Monkey P's variability instead increases with reward, and Monkey R's shows little modulation.

(E) Peak speed variability generally decreases with greater reward. Both Monkeys E and R show less variability in reaction time with greater reward, while Monkey P shows no visible difference.

From this analysis, we conclude that reward may modulate variability in our constituent metrics of vigor, though does not necessarily do so in a consistent way across animals. These intersubject differences may pertain to differences in how the subjects process reward-mediated strategy changes (see Discussion).

Finally, we also want to note that this task was challenging, where target size and reach time limit were carefully titrated for each animal to push their performance to their limits.

(F) Animals succeeded at the task ~75% of the time with a variety of failure modes accounting for the remainder of trials. We categorized each trial as either a success, or one of 5 possible failure modes (Methods)[75]. Arranged from the top of the bar being roughly earlier in the trial to bottom being success: 1. False Start (dark red), 2. Delay Drift (red), 3. Overshoot (light red), 4. Undershoot (orange), 5. Target Hold Fail (olive), or Success (green).

While we do not observe differences in our findings form the neural data if we include versus exclude failure trials, we do note that a sufficiently challenging task may be necessary to observing reward-related changes in behavior or neural activity[88].

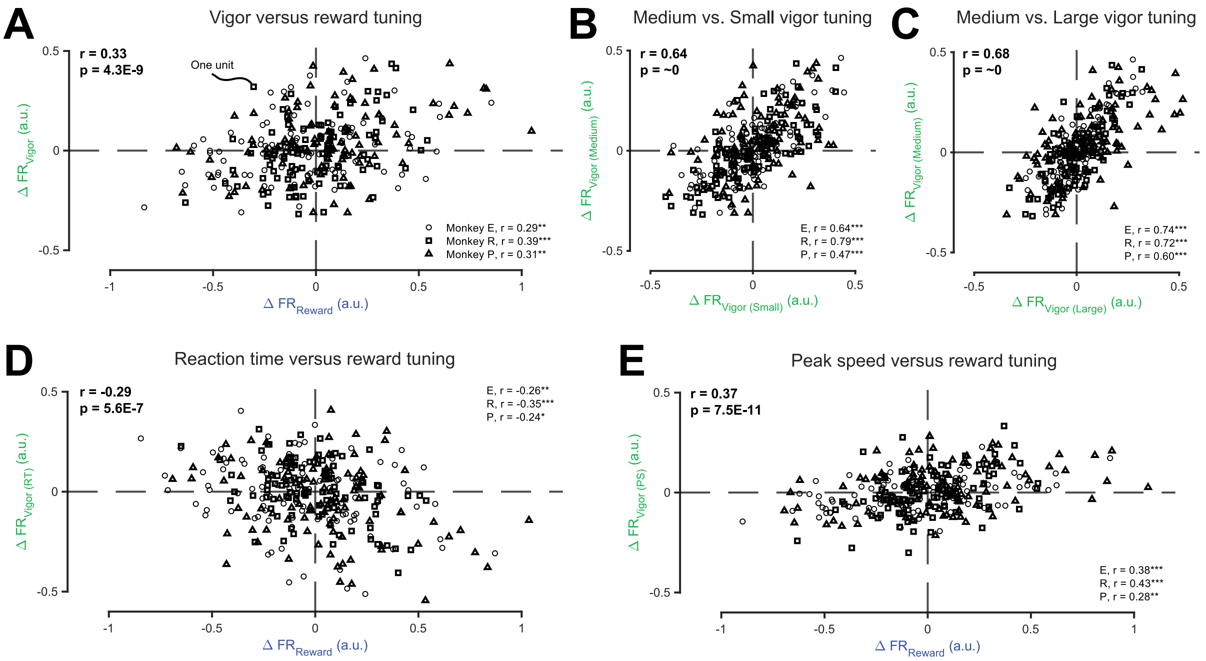


**Figure S2. Comparing single unit vigor and reward tuning during movement preparation.**

We wanted to assess if single unit tuning to reward and vigor was correlated during movement preparation (e.g., units with higher firing rate for Large reward also fire more for upcoming High vigor movements). Because the recordings were using chronically implanted electrode arrays, there was considerable overlap in units between days. This means we cannot assess neural units as independent on different sessions. We leveraged the results of the chronic unit identification algorithm described above for use in neural stitching (Methods) to identify consistent units. For this analysis, we only kept the 100 units from each animal that were present for the greatest number of trials. We did this for three reasons: (1) To only use units with many trials to gain statistical power, (2) To reduce the likelihood of "double-counting" units that were improperly identified as different across sessions, and (3) To use an equal number of units for each animal in overall statistics. Keeping 50 or 200 units instead produced near-identical results and does not alter interpretations.

We calculated single unit firing rates from a bin of activity at the end of movement preparation (from 150 ms before go cue to 50 ms after). For each unit, we z-scored firing rates and vigor within each day-direction condition over its available trials. To calculate reward tuning, we first controlled for vigor by repeatedly subsampling trials to match vigor distributions for Small and Large reward trials (see **Fig. 8A,** Methods). We then took the average firing rate for Small and Large vigor-matched subsamples and calculated the difference between rewards ($\Delta FR_{Reward}$ ; positive values indicate greater firing rate for Large rewards). Similarly, on Medium reward trials, we calculated the average firing rate for trials where movements were Low and High vigor (i.e., below or above the median vigor) and calculated the difference between them ($\Delta FR_{Vigor}$ ; positive values indicate greater firing rate for High vigor).

(A) Motor cortical tuning to cued reward and upcoming movement vigor are correlated during movement preparation. We show $\Delta FR_{Reward}$ versus $\Delta FR_{Vigor}$ for the 100 units present for the most trials/sessions for each animal (Methods). We then plotted these against each other and calculated Spearman rank correlation (*$p < 0.05$, **$p < 0.01$, ***$p < 0.001$, correlation t-test). Overall results (upper-left) and single animal (bottom-right) are shown. Axis units are on the same scale as one another, as both are firing rates z-scored using all trials' variance.

Along with using Medium reward trials as described above ($\Delta FR_{Vigor}$, or $\Delta FR_{Vigor\ (Medium)}$), we calculated vigor tuning within Small ($\Delta FR_{Vigor\ (Small)}$) and Large ($\Delta FR_{Vigor\ (Large)}$) trials as well to assess the consistency of vigor tuning across reward conditions. Low versus High vigor labels were assigned based on the median vigor separately for each reward condition.

(B-C) Vigor tuning is highly correlated between different reward conditions. Same format as panel A. Panel B shows Small reward trial vigor tuning along the x-axis, and panel C shows Large. For reference, we note that when we calculated correlations in vigor tuning between subsets of trials within a given reward condition, values ranged from 0.5-0.8. “p = ~0” indicates p-value is within machine-precision of 0.

(D-E) Finally, we also compared vigor with reward tuning separately using the constituent metrics of vigor, reaction time (panel D) and peak speed (panel E). Same format as panel A. Results match those of the combined vigor metric.

These results support three conclusions about the nature of reward and vigor tuning at the end of movement preparation. First, reward and vigor tuning are significantly correlated with one another (panel A). Second, vigor tuning is consistent across reward conditions (panel B-C). Finally, reward tuning is correlated with the individual components of vigor as well (panels D-E).

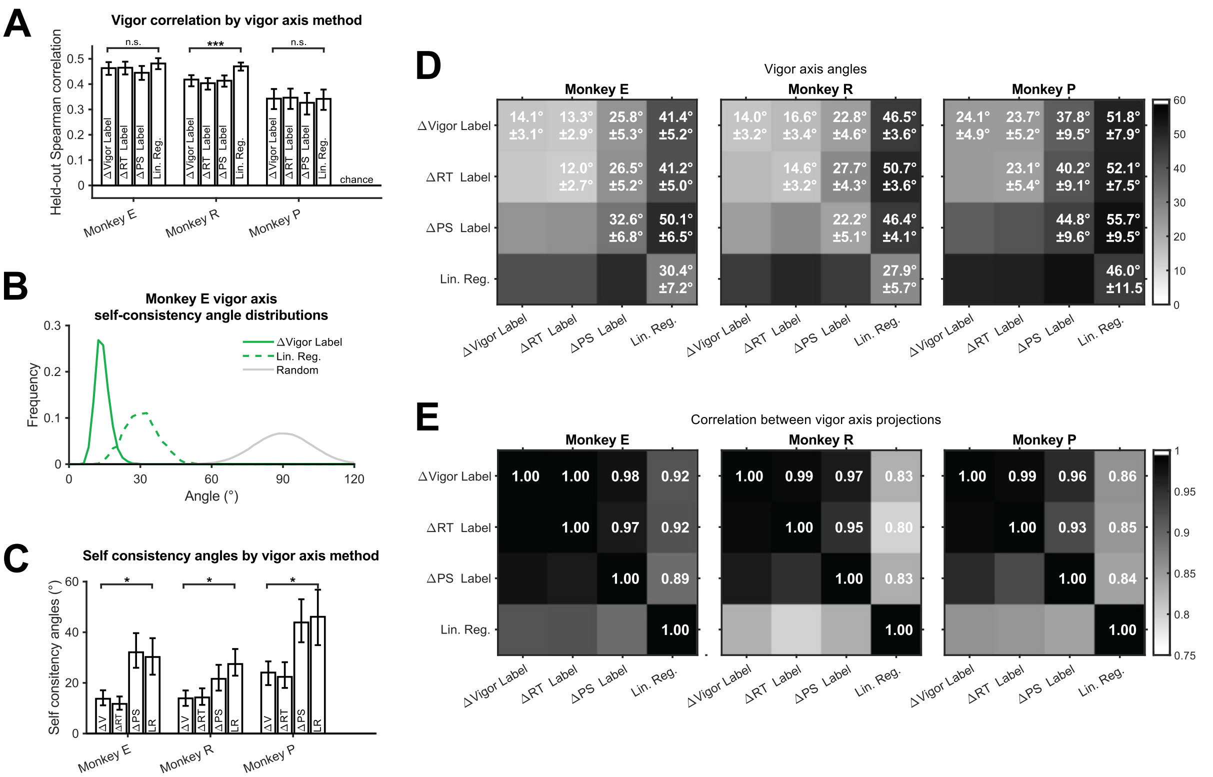


**Figure S3. Comparison of vigor axis methods.**

There are a variety of ways one can find a dimension of neural activity that relates to movement vigor. In the main text, we identified a vigor axis in the neural preparatory activity state space by finding the vector that connected the average activity for Low vigor label trials with the average for High vigor trials within Medium rewards (**Fig. 3C**), which for this figure we will call the *ΔVigor Label* method. We tested three other methods for identifying a vigor axis. We performed the same procedure but using reaction time (*ΔRT Label* method) or peak speed (*ΔPS Label* method) as the labels instead. Also, we identified a vigor axis using linear regression (*Lin. Reg.* method) of neural activity to trial-by-trial vigor values. We used principal components regression and selected the number of components to maximize held-out data correlation with vigor (Monkey E = 11 components, R = 18, P = 8). We performed this analysis on the stitched neural data, which effectively performs dimensionality reduction via factor analysis while combining data across days (Methods), and mean-centered neural activity and vigor within each day-direction condition.

For each method, we wanted to assess (1) How well projections along the vigor axis correlated with vigor, (2) How consistent estimation of the vigor axis was, and (3) How similar the axes were to one another. To accomplish this, we repeatedly (2000 times) randomly split our trials in half between “training” and “test” sets, calculated the vigor axes in both, and projected the test set trials along the vigor axes found in the training set. Only Medium trials were used for this analysis.

(A) vigor-axis projections show similar levels of correlation with vigor across methods. We calculated the correlation between test set vigor-axis projections and movement vigor values and compared results between vigor axis methods. We show the median correlation (with 68.2% confidence interval, akin to standard error) between vigor-axis projections and vigor values in test data. We performed statistical comparisons between the *ΔVigor Label* and *Lin. Reg.* methods by comparing values within each repeat’s test set, where a p-value was calculated using the ratio of values in one greater/less than the other; for example, since the test is two-sided, a situation where a value in one condition was greater than the other 2.5% of the time would correspond to $p = 0.05$ (***$p < 0.001$, n.s. = not significant). Only Monkey R showed a significant improvement using *Lin. Reg.* compared to *ΔVigor Label*, and the improvement was small (correlation difference < 0.05).

To determine how reliably we could estimate a given vigor axis, we calculated “self consistency angles” as the angles between vigor axes found from the training and test sets within each method. The more reliably an axis can be estimated, the closer the distribution of values over repeats is to 0 degrees.

(B) For visualization, we show distributions of self consistency angles over repeated subsamples for one animal (Monkey E) for the *ΔVigor Label* and *Lin. Reg.* methods. We also show a distribution of angles between random vectors in the stitched neural space (Monkey E: 21 factors).

(C) Vigor axes are estimated more reliably using the *ΔVigor Label* method than the *Lin. Reg.* method. Same format and method of statistical comparison as panel A but using self consistency angles instead of correlation with vigor (*$p < 0.05$). For all animals, the self consistency angles were significantly lower for the *ΔVigor Label* method than for the *Lin. Reg.* method, typically by approximately a factor of 2.

From these results so far, we conclude that the *ΔVigor Label* method used in the main text identifies vigor axes more reliably than the *Lin. Reg.* method with little-to-no detriment to correlations with behavior. We note that the animal that did show a significant increase in correlation for the *Lin. Reg.* method also has by far the most data (Monkey R has ~2x Monkey E’s number of trials and ~8x Monkey P’s).

We lastly wanted to compare the axes to one another to determine how similar or different of vigor axes the methods produced. We did this by evaluating two quantities. First, we calculated the angle between the training set vigor axis found using one method with the test set vigor axis using the others. Second, we calculated the correlation between the vigor axes’ projections using all trials. We would expect that similar axes have an angle that is close to a given method’s self-consistency angle and that they would show high correlation of projections with one another.

(D) Different methods produce fairly aligned vigor axes. We show the mean (± S.E., see Methods) of the angle between the vigor axes found in the training sets and test sets. The diagonal elements show self consistency angles. We generally find that the *ΔLabel* methods produce axes that are well-aligned with one another, with the angle between any pair near-matching the greater of the two’s self consistency angles. The *Lin. Reg.* method produced axes not perfectly aligned with the other methods, but much closer to aligned than orthogonal.

(E) Correlation of vigor-axis projections found with different methods are well-correlated. We calculated the Spearman rank correlation between vigor-axis projections in the test sets for each subsample and show the average. vigor-axis projections found using the *ΔLabel* methods were near-perfectly correlated with one another (average = 0.97), and were still very positively correlated with projections found using the *Lin. Reg.* method (average = 0.86).

These results indicate that all of the *ΔLabel* methods are effectively estimating the same vigor axis. This implies that encoding of vigor metrics from preparatory neural activity state is uni-dimensional, as opposed to reaction time and peak speed being encoded distinctly from one another.

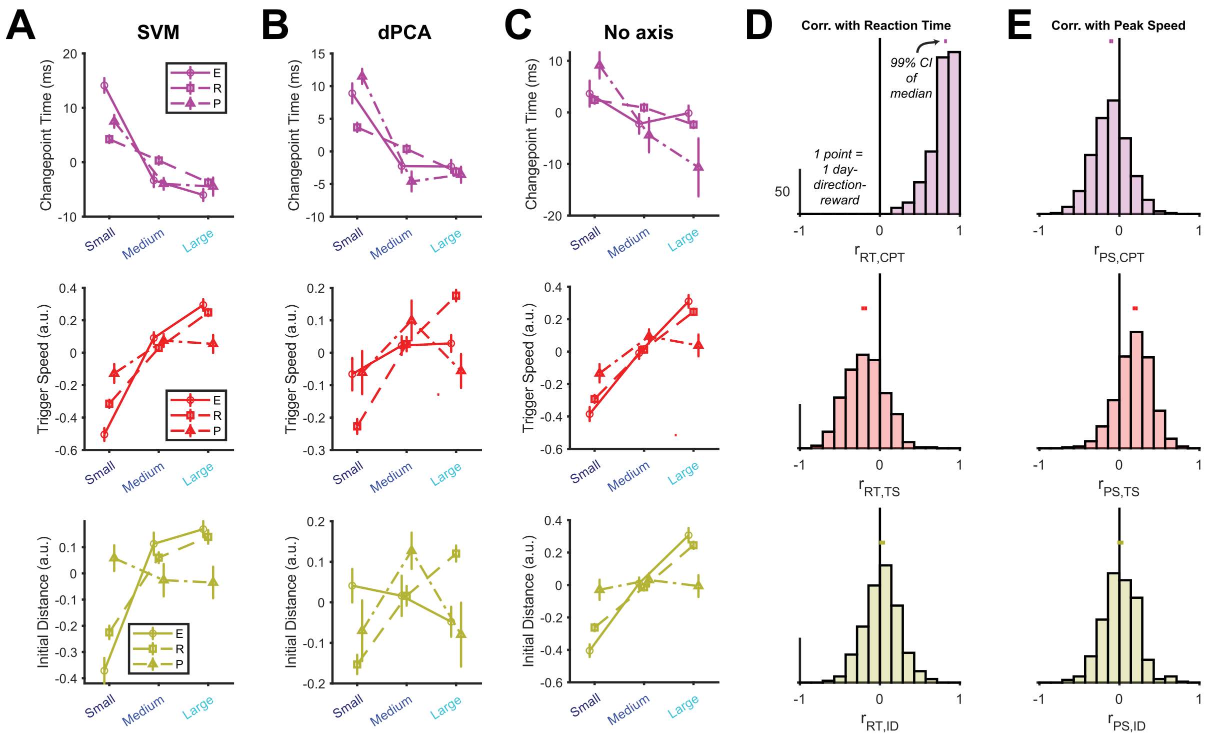


**Figure S4. Alternative methods for trigger axis metrics.**

We tested other methods of identifying the trigger axis and calculating the corresponding metrics shown in **Figure 5**. We only show reward effects here for space, but all significant correlations with vigor metrics that we observed with these metrics found using an LDA-based trigger axis (**Fig. 5**) hold for each of these methods as well.

(A) SVM: We used a linear support vector machine classifier to identify a maximum-margin dimension separating the same pre- and peri-movement bins of neural activity as used with LDA in **Fig. 5** (regularization parameter $\lambda = (2n)^{-1}$, where n is number of trials). This is similar to the method used in [68]. Results are near-identical to LDA's.

(B) dPCA: We performed demixed principal components analysis to demix direction, reward, and condition-invariant information from 250 ms before movement onset to 250 ms after, then dubbed the highest-variance condition-invariant dimension as the trigger axis. This is similar to the method used in [36,38]. The results are generally similar to LDA and SVM.

(C) No axis: instead of finding a trigger axis, we calculated neural speed in the full factor space of the stitched neural data. We then treated this neural speed as the trigger axis speed for calculating trigger speed and changepoint time (Methods). For initial distance, we calculated the euclidean distance in the full factor space on each trial between the state at changepoint time and the state at movement onset. As such, this approach calculates all of the same metrics as previous, but does so without ever actually identifying a trigger axis. Results were quite similar to the LDA approach.

We additionally considered the correlations of trigger axis metrics found using LDA with reaction time and peak speed separately, as opposed to with the combined vigor metric (main text).

(D) Reaction time correlations with trigger axis metrics. Same format as **Fig. 5D,F,H**.

(E) Peak speed correlations with trigger axis metrics.

We note that the trends in correlations for all three trigger axis metrics for both reaction time and peak speed mirror that in vigor overall. That is, faster changepoint times and trigger speeds are significantly correlated with greater vigor (lower reaction time and higher peak speed), but that initial distance shows little correlation with behavior. However, the extent of correlation is not necessarily the same; in particular, changepoint time is very highly correlated with reaction time, but only weakly with peak speed.

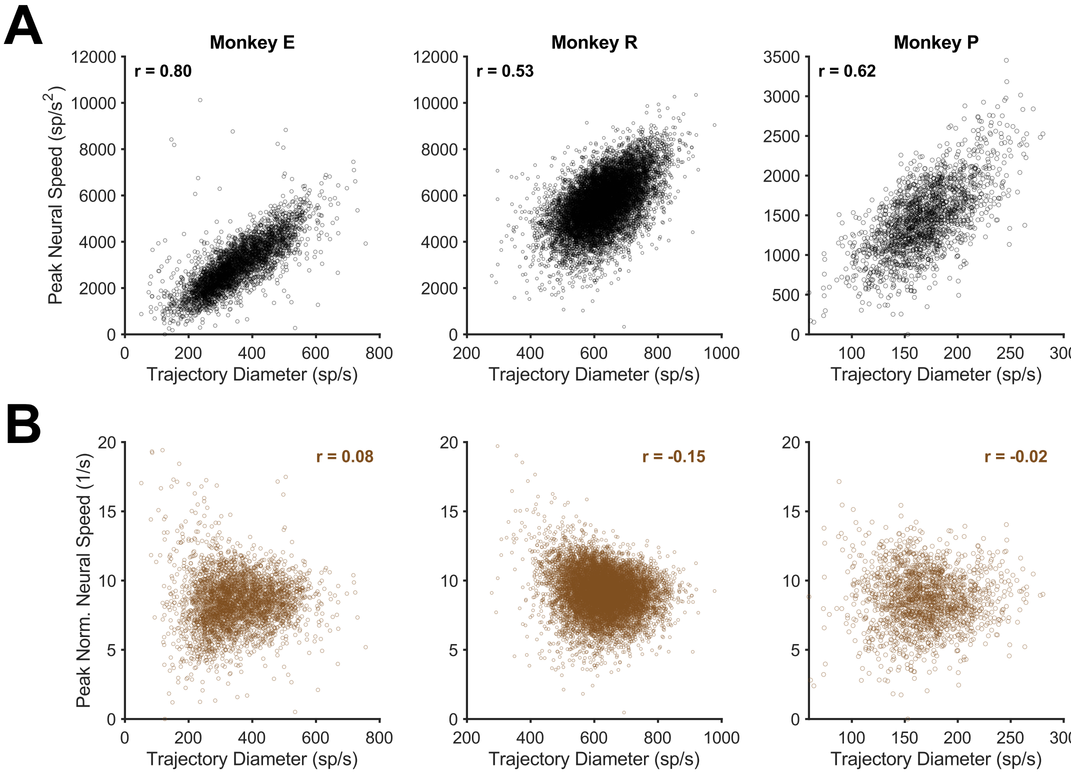


**Figure S5. Normalizing peak neural speed by the trajectory diameter decorrelates the two quantities.**

When assessing how reward and vigor affect neural trajectories during movement execution, we wanted to examine both the magnitude of how far the trajectory stretched each trial (trajectory diameter) and also how quickly the neural activity proceeded along the trajectory (neural speed). While trajectory diameter calculation was fairly straightforward (**Fig. 6C**), calculating a speed along the trajectory was nontrivial. Our solution was to calculate a peak neural speed (i.e., maximum of time derivative of trajectory), then normalize by dividing each trial's value by its trajectory diameter (Methods). We examined this intermediate peak neural speed quantity and compared it to trajectory diameter.

(A) Peak neural speed is highly correlated with trajectory diameter. We show the peak neural speed (*not* normalized) from each trial scattered against the trajectory diameter for each animal and show the Spearman rank correlation in insets at the top-left. For sake of evaluation, we did not perform any z-scoring within day-direction conditions for this figure. This relationship between peak neural speed and trajectory diameter is intuitive and not dissimilar to phenomena seen relating values and their time derivatives in natural kinematics (e.g., the "main sequence" relating saccade distance and velocity).

(B) After normalizing by trajectory diameter, correlations between neural speed and trajectory diameter are effectively eliminated. We scattered the peak normalized neural speed versus the trajectory diameter and show Spearman rank correlation in insets in the top-right corner.

As such, we have scientific and empirical reason to normalize neural speed by the trajectory diameter. Scientifically, the quantity we want to approximate is a speed along the canonical trajectory shape, which should scale with the size of the trajectory and hence should account for it. Empirically, using raw peak neural speed results in a metric that is highly correlated with trajectory diameter and, as such, providing little additional information; conversely, the peak normalized neural speed shows only weak (if any) correlation with trajectory diameter.

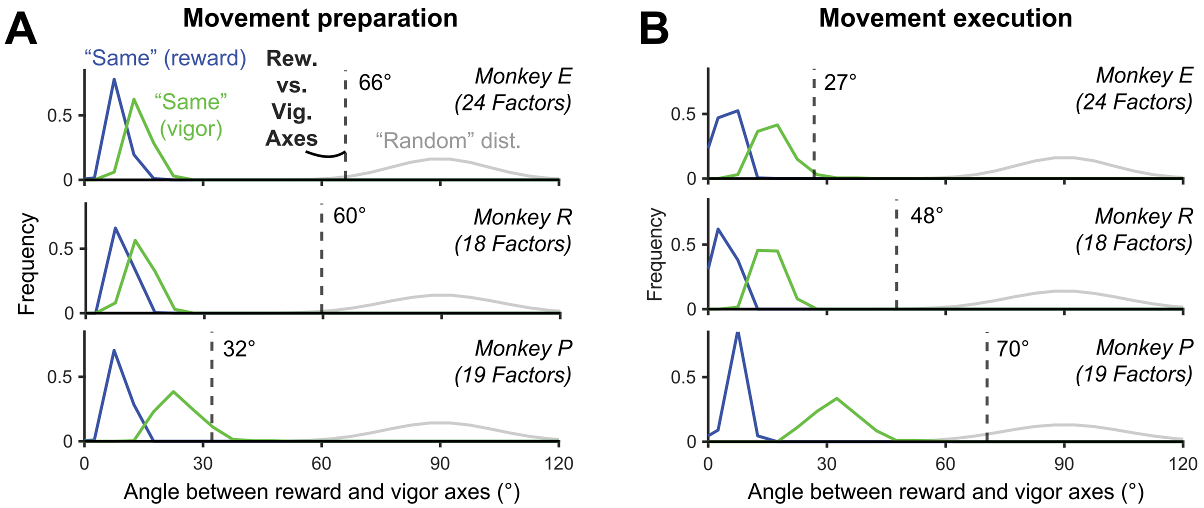


**Figure S6. Reward and vigor axes are neither orthogonal nor well-aligned.**

We wanted to assess if the vigor and reward axes within a given epoch were orthogonal, well-aligned, or somewhere in between. To do this, we repeated the following procedure 1000 times: we randomly chose half of the trials from each reward condition to be "set 1" and the other half to be "set 2". Within each set, we calculated both a vigor and reward axis for each epoch (see Methods for axis calculation). To assess how reliably we could estimate a given axis, we calculated the angle between the same axis found in set 1 and set 2, which we call the "self-consistency angle" elsewhere (**Fig. S3**). We show the distributions of these self-consistency angles across the 1000 random set splits for each animal and epoch as "Same" distributions (blue for reward axis green for vigor axis).

To assess the average angle between the reward and vigor axes, we calculated the angle between set 1 reward axis and set 2 vigor axis, along with set 1 vigor axis and set 2 reward axis, then took the median of these values across all splits (total of 2000 values; black dashed line). Finally, we also formed a random vector distribution in the neural space by drawing random vectors from an isotropic multivariate Gaussian distribution and calculating their angles with one another (100000 vectors; gray). Overall, this permits us to compare the angle between reward and vigor axes with self-consistency angles and what would be predicted from random axes.

(A) During movement preparation, the reward and vigor axes are neither consistently well-aligned nor orthogonal to each other.

(B) During movement execution, the reward and vigor axes are neither consistently well-aligned nor orthogonal to each other.

While we observe some potentially interesting trends between the different animals, we overall find that the angle between the reward and vigor axes tends to land somewhere between the top percentiles of the self-consistency angles and the bottom of the random vector distribution.

**Table S1. All main figure statistics.** ~0 indicates values within machine precision of 0.

| Figure | Subject | n | Quantification | Value | Test | P-value | Shown |
|---|---|---|---|---|---|---|---|
| **1B** | E | S = 1112 trials, M = 1348, L = 1320 | Cursor speed | - | - | - | Mean +/- S.E. shading |
| **1C** | E | Same as 1B | Large trial median reaction time minus Small trial median | - | Wilcoxson rank sum | 3.9E-31 | Median +/- S.E. bars |
| | | | Large trial mean peak speed minus Small trial mean | - | Welch's t-test | 8.4E-29 | Mean +/- S.E. bars |
| | R | S = 3133 trials, M = 3364, L = 3494 | Large trial median reaction time minus Small trial median | - | Wilcoxson rank sum | 2.6E-13 | Median +/- S.E. bars |
| | | | Large trial mean peak speed minus Small trial mean | - | Welch's t-test | 4.9E-17 | Mean +/- S.E. bars |
| | P | S = 616, M = 682, L = 584 | Large trial median reaction time minus Small trial median | - | Wilcoxson rank sum | 7.2E-15 | Median +/- S.E. bars |
| | | | Large trial mean peak speed minus Small trial mean | - | Welch's t-test | 0.002 | Mean +/- S.E. bars |
| **1D** | All | 162 day-direction conditions | Median of Large minus Small average vigor | 0.31 | Sign rank test | 2.4E-19 | All data points |
| | E | 48 | | 0.46 | | 7.3E-9 | |
| | R | 94 | | 0.27 | | 1.8E-9 | |
| | P | 20 | | 0.32 | | 0.002 | |
| **2A** | E (unit 299) | S = 363 trials, L = 440 | Firing rate | - | - | - | Mean +/- S.E. shading |
| | E (unit 66) | S = 1112, L = 1320 | | - | - | - | |
| | E (unit 106) | S = 1112, L = 1320 | | - | - | - | |
| **2B** | E (unit 299) | Low = 179 trials, High = 340 | Firing rate | - | - | - | Mean +/- S.E. shading |
| | E (unit 66) | Low = 600, High = 748 | | - | - | - | |
| | E (unit 106) | Low = 600, High = 748 | | - | - | - | |
| **2D** | E | S = 1047 trials, M = 1291, L = 1269 | Large trial average vigor axis projection minus Small average | - | Welch's t-test | 1.6E-65 | Mean +/- S.E. bars |
| | R | S = 2264, M = 2472, L = 2509 | | - | | 1.2E-63 | |
| | P | S = 451, M = 509, L = 396 | | - | | 1.2E-82 | |
| **2E** | All | 318 day-direction-reward conditions (Small and Large) | Median correlation of vigor with vigor axis projections across conditions | 0.35 | Sign rank test | 3.8E-49 | All data points |
| | E | 96 | | 0.38 | | 5.7E-16 | |
| | R | 182 | | 0.35 | | 3.2E-29 | |
| | P | 40 | | 0.36 | | 3.5E-7 | |
| **3C** | E | S = 1047 trials, L = 1269 | Average projection in target plane | - | - | - | - |
| **3D** | E | Low = 634 trials, High = 635 | Average projection in target plane | - | - | - | - |
| **3F** | E | Same as 2D | Large trial average target ring radius minus Small average | - | Welch's t-test | 7.0E-41 | Mean +/- S.E. bars |
| | R | | | - | | 1.1E-20 | |
| | P | | | - | | 1.4E-24 | |

| | | | | | | | |
|---|---|---|---|---|---|---|---|
| **3G** | All | 476 day-direction-reward conditions | Median correlation of vigor with vigor axis projections across conditions | 0.16 | Sign rank test | 1.7E-32 | All data points |
| | E | 144 | | 0.22 | | 1.9E-14 | |
| | R | 272 | | 0.15 | | 1.5E-15 | |
| | P | 60 | | 0.15 | | 6.6E-6 | |
| **4C** | E | 3608 trials | Correlation of reaction time with trigger time | 0.85 | - | - | All data points |
| | R | 9883 | | 0.93 | | - | |
| | P | 1850 | | 0.76 | | - | |
| **4D** | E | S = 1049 trials, M = 1285, L = 1274 | Large trial median trigger time minus Small median | - | Wilcoxson rank sum | 1.1E-64 | Median +/- S.E. bars |
| | R | S = 3096, M = 3328, L = 3459 | | - | | 3.2E-18 | |
| | P | S = 615, M = 676, L = 559 | | - | | 4.9E-25 | |
| **5C** | E | Same as 4D | Large trial median changepoint time minus Small median | - | Wilcoxson rank sum | 1.8E-46 | Median +/- S.E. bars |
| | R | | | - | | 3.4E-20 | |
| | P | | | - | | 1.4E-14 | |
| **5D** | All | 489 day-direction-reward conditions | Correlation of vigor with changepoint time | -0.61 | Sign rank test | 8.4E-82 | All data points |
| | E | 144 | | -0.61 | | 2.3E-25 | |
| | R | 285 | | -0.64 | | 1.7E-48 | |
| | P | 60 | | -0.40 | | 2.2E-11 | |
| **5E** | E | Same as 4D | Large trial mean trigger speed minus Small mean | - | Welch's t-test | 1.2E-119 | Mean +/- S.E. bars |
| | R | | | - | | 3.0E-98 | |
| | P | | | - | | 1.1E-5 | |
| **5F** | All | Same as 5D | Correlation of vigor with trigger speed | 0.23 | Sign rank test | 2.4E-43 | All data points |
| | E | | | 0.16 | | 8.6E-12 | |
| | R | | | 0.26 | | 1.0E-35 | |
| | P | | | 0.32 | | 8.0E-11 | |
| **5G** | E | Same as 4D | Large trial mean initial distance minus Small mean | - | Welch's t-test | 7.2E-19 | Mean +/- S.E. bars |
| | R | | | - | | 4.1E-44 | |
| | P | | | - | | 5.0E-3 | |
| **5H** | All | Same as 5D | Correlation of vigor with initial distance | -0.00 | Sign rank test | 0.23 | All data points |
| | E | | | -0.01 | | 0.30 | |
| | R | | | 0.02 | | 0.76 | |
| | P | | | -0.05 | | 0.25 | |
| **6D** | E | S = 970 trials, M = 1268, L = 1276 | Large trial median trajectory diameter minus Small median | - | Wilcoxson rank sum | 7.0E-115 | Median +/- S.E. bars |
| | R | S = 3126, M = 3359, L = 3494 | | - | | 5.6E-148 | |
| | P | S = 610, M = 673, L = 570 | | - | | 0.47 | |
| **6E** | All | 489 day-direction-reward conditions | Correlation of peak speed with trajectory diameter | 0.19 | Sign rank test | 3.3E-54 | All data points |
| | E | 144 | | 0.23 | | 2.4E-18 | |
| | R | 285 | | 0.19 | | 4.1E-34 | |
| | P | 60 | | 0.15 | | 3.5E-5 | |
| **6G** | E | Same as 6D | Large median peak normalized neural speed minus Small median | - | Wilcoxson rank sum | 6.2E-22 | Median +/- S.E. bars |
| | R | | | - | | 8.5E-30 | |
| | P | | | - | | 2.6E-7 | |
| **6H** | All | Same as 6E | Correlation of peak speed with peak normalized neural speed | 0.10 | Sign rank test | 1.9E-23 | All data points |
| | E | | | 0.06 | | 3.2E-3 | |
| | R | | | 0.14 | | 3.4E-21 | |
| | P | | | 0.08 | | 2.4E-3 | |

| | | | | | | | |
|---|---|---|---|---|---|---|---|
| **7B** | E | 1197 Medium trials with values for all neural metrics | Correlation between all neural metrics identified | - | - | - | All correlations |
| | R | 2443 | | - | - | - | |
| | P | 488 | | - | - | - | |
| **8B** | E | S and L = 761 trials in each subsample | Unmatched distribution average decoding accuracy minus matched distribution's | - | Modified t-test (see Methods) | See figure | Mean across folds/reps +/- average S.E. of folds over reps |
| | R | S and L = 2780 | | - | | | |
| | P | S and L = 449 | | - | | | |
| **8D** | E | Same as 8B | Average percentage of Small to Large neural distance for matched distribution compared to unmatched | - | - | - | Mean across reps |
| | R | | | - | - | - | |
| | P | | | - | - | - | |
| **8F** | All | 324 day-direction-epoch conditions | Median difference in Small-to-Large effect size (Cohen's d) between vigor axis and orth. reward axis projections | 0.68 | Sign rank test | 2.8E-41 | All data points |
| | E | 96 | | 0.69 | | 5.3E-11 | |
| | R | 188 | | 0.67 | | 1.1E-28 | |
| | P | 40 | | 0.72 | | 3.5E-5 | |
